# Human pluripotent stem cell-derived kidney organoids for personalized congenital and idiopathic nephrotic syndrome modeling

**DOI:** 10.1101/2021.10.27.466054

**Authors:** Jitske Jansen, Bartholomeus T van den Berge, Martijn van den Broek, Rutger J Maas, Brigith Willemsen, Christoph Kuppe, Katharina C Reimer, Gianluca Di Giovanni, Fieke Mooren, Quincy Nlandu, Helmer Mudde, Roy Wetzels, Dirk den Braanker, Naomi Parr, James S Nagai, Vedran Drenic, Ivan G Costa, Eric Steenbergen, Tom Nijenhuis, Nicole Endlich, Nicole CAJ van de Kar, Rebekka K Schneider, Jack FM Wetzels, Johan van der Vlag, Rafael Kramann, Michiel F Schreuder, Bart Smeets

**Author notes:** These authors contributed equally (Bartholomeus T van den Berge, Martijn van den Broek, Rutger J Maas). These authors contributed equally (Michiel F Schreuder, Bart Smeets). <u>Corresponding authors:</u> Bart Smeets, PhD, Jitske Jansen, PhD, Radboud University Medical Center, Radboud Institute for Molecular Life Sciences, Department of Pathology, P.O. Box 9101, 6500 HB Nijmegen, the Netherlands.

## Abstract

Nephrotic syndrome (NS) is characterized by severe proteinuria as a consequence of kidney glomerular injury due to podocyte damage. *In vitro* models mimicking *in vivo* podocyte characteristics are a prerequisite to resolve NS pathogenesis. Here, we report human induced pluripotent stem cell derived kidney organoids containing a podocyte population that heads towards adult podocytes and were superior compared to 2D counterparts, based on scRNA sequencing, super-resolution imaging and electron microscopy. In this study, these next-generation podocytes in kidney organoids enabled personalized idiopathic nephrotic syndrome modeling as shown by activated slit diaphragm signaling and podocyte injury following protamine sulfate treatment and exposure to NS plasma containing pathogenic permeability factors. Organoids cultured from cells of a patient with heterozygous *NPHS2* mutations showed poor *NPHS2* expression and aberrant *NPHS1* localization, which was reversible after genetic correction. Repaired organoids displayed increased VEGFA pathway activity and transcription factor activity known to be essential for podocyte physiology, as shown by RNA sequencing. This study shows that organoids are the preferred model of choice to study idiopathic and congenital podocytopathies.

**Summary Statement:** Kidney organoid podocytes allow personalized nephrotic sydrome modeling,

## Introduction

Podocytes play an important role in the glomerular filtration barrier. Podocytes possess interdigitating foot processes that are bridged by a protein complex called the slit diaphragm, which contains proteins such as nephrin (*NPHS1*) and podocin (*NPHS2*). Malfunctioning of podocytes (podocytopathy) causes massive proteinuria, resulting in the nephrotic syndrome (NS) (Shabaka et al., 2020; Veissi et al., 2020). In patients with idiopathic NS (iNS) a kidney biopsy shows no abnormalities in light microscopy. Complete foot process effacement, as determined by electron microscopy, defines iNS as a podocytopathy. The identification of mutations in congenital nephrotic syndrome in important podocyte genes (*NPHS1* and *NPHS2*) (Maas et al., 2016; Noone et al., 2018) also defines iNS as a podocytopathy. The pathophysiology of idiopathic NS has not been clarified yet. Within the spectrum of NS, a unique subset of patients have presumed pathogenic circulating permeability factors (CPFs) that result in podocyte injury and eventually recurrence of the disease after kidney transplantation. Clinical evidence suggests CPFs involve the immune system (i.e. T- and B-cells) (Colucci et al., 2018; Gallon et al., 2012; Straatmann et al., 2014). However, there is still no consensus about the nature of the CPFs (Maas et al., 2014; Rudnicki, 2016; Shoji et al., 2020).

In order to gain more insight in podocytopathies, animal models and basic *in vitro* models have been explored to gain more insight in podocytopathies, but these failed to provide full mechanistic insight. Hence, there is an unmet need for models which recapitulate glomerular physiology and pathology to resolve the molecular mechanism of underlying glomerular diseases. Previously, podocyte cell lines, human induced pluripotent stem cell (iPSC)-derived 2D podocytes, and 3D kidney organoid models have been established (Garreta et al., 2019; Musah et al., 2017; Rauch et al., 2018; Saleem et al., 2002; Sharmin et al., 2016; Takasato et al., 2015; Wu et al., 2018). The advantages of 2D podocyte models are the straightforward and standardized culture conditions, though the presence and localization of slit diaphragm proteins are not convincing (Hale et al., 2018; van den Broek et al., 2021). Kidney organoid models are more complex and require challenging culture protocols, but podocyte characteristics, including slit diaphragm proteins, have been shown (Hale et al., 2018; Taguchi and Nishinakamura, 2017; van den Berg et al., 2018; Wu et al., 2018). Despite the substantial progress that has been made in the kidney organoid field, current organoids resemble immature tissue, show limited vascularization, and do not have a functional glomerular filtration barrier (Takasato et al., 2015; Uchimura et al., 2020; van den Berg et al., 2018). Recently, Tanigawa and colleagues successfully modeled Finnish-type congenital NS (*NPHS1* mutant) in kidney organoids, identified slit diaphragm abnormalities in these podocytes and could genetically correct the mutation (Tanigawa et al., 2018). The study by Tanigawa *et al*. emphasized the potential of kidney organoids for personalized regenerative medicine and to accurately study NS *in vitro*.

The aim of our study was to investigate the potency of current kidney organoids to accurately model podocytopathies, as observed in NS. Here, we established a hybrid differentiation protocol that resulted in kidney organoids containing a podocyte population that heads towards adult podocyte expression levels. We succeeded in congenital (*NPHS2* mutant) NS modelling as well as idiopathic NS modeling. Our data here indicate that iPSC derived kidney organoids are the preferred system to model NS *in vitro*, since cellular signaling cascades are induced upon injury that are absent in 2D iPSC-derived podocytes and a podocyte cell line.

## Results

### Podocytes in kidney organoids show superior characteristics compared to 2D podocytes

Human iPSCs were successfully cultured into kidney organoids using a hybrid directed differentiation protocol based on Uchimura and Takasato et al (Takasato et al., 2015; Uchimura et al., 2020). Using this approach, segmented patterning was induced and organoids consisted of kidney cells including podocytes, proximal tubule, loop of Henle, distal tubule, and collecting duct-like cells, as well as endothelial and mesangial cell precursors, stromal cells, and off-target neuronal progenitors, as shown by single-cell RNA sequencing and immunofluorescence staining (Figure 1B – C, Figure S1, Supplemental Table S1).

**Figure 1.**
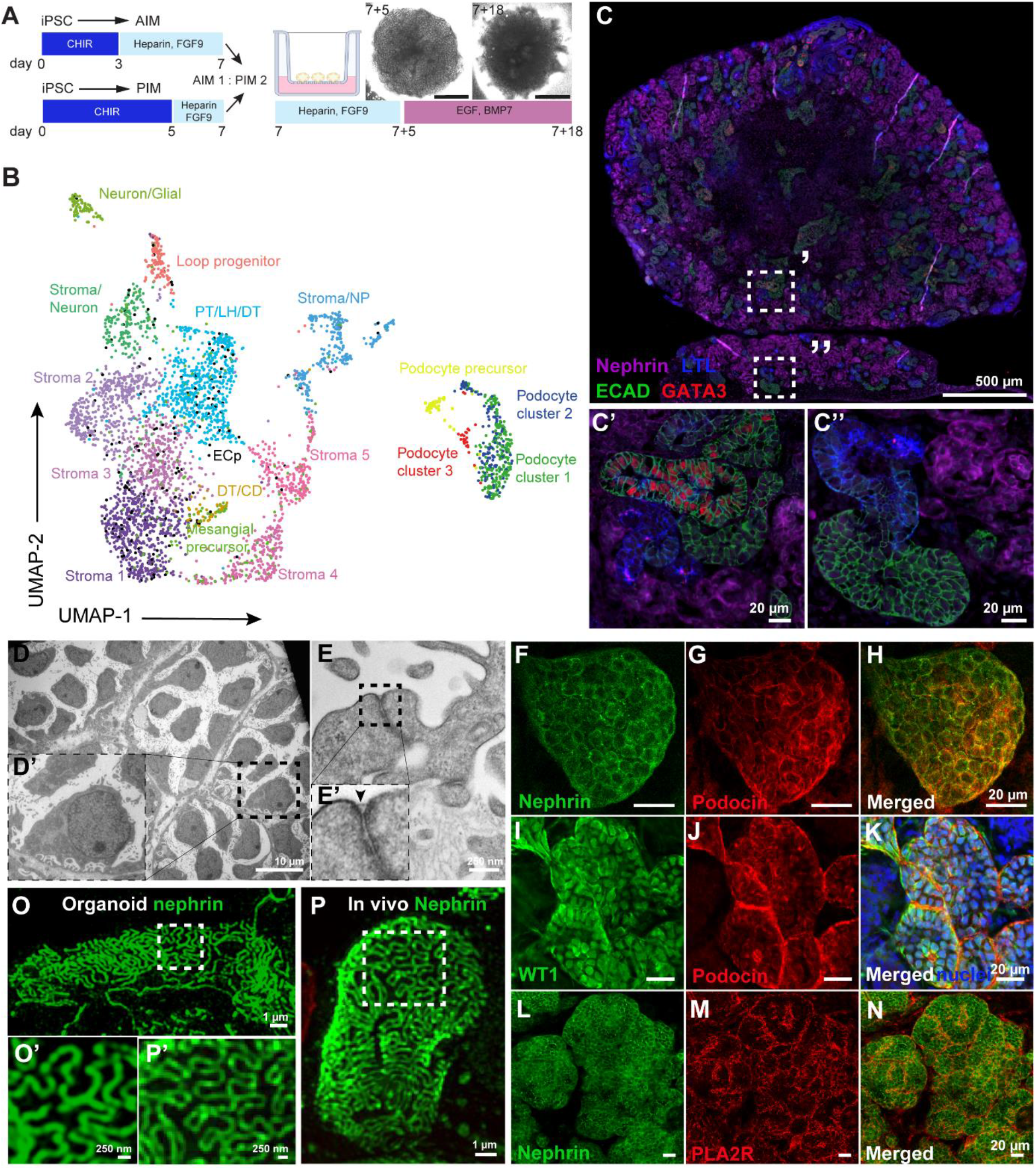
Human induced pluripotent stem cell (iPSC)- derived kidney organoids contain 17 distinct cell clusters including well-developed podocytes showing filtration slits. (A) Schematic representation of the iPSC-derived kidney organoids culture. iPSCs were differentiated in 2D either towards anterior intermediate mesoderm (AIM) or posterior intermediate mesoderm (PIM) and aggregates (300,000 cells) were made using a 1:2 AIM/PIM ratio on day 7. Organoids were cultured on the air-liquid interface of Transwell™ filters for an additional 18 days, using a mixture of growth factors as indicated in Figure 1A. C: CHIR99021, F9: Fibroblast growth factor 9, EGF: epidermal growth factor, BMP7: bone morphogenic protein 7. (B) Uniform Manifold Approximation and Projection (UMAP) integration of 3,900 single cells of human iPSC-derived kidney organoids results in 17 different cell clusters. PT: proximal tubular cells, LH: Loop of Henle cells, DT: distal tubular cells, NP: neuronal progenitors, CD: collecting duct precursors, ECp: endothelial cell precursors. (C-C’’) Segmented patterning in human kidney organoids shows the presence of podocytes (NPHS1, magenta), proximal tubules (Lotus Tetragonolobus Lectin (LTL), blue), distal-like (E-cadherin (ECAD), green) and collecting duct-like cells (ECAD (green) and GATA binding protein 3 (GATA3) (red). (D-D’) Ultrastructural analysis of organoid-derived podocytes showing cell bodies and interdigitating foot processes including (E-E’) a filtration slit, indicated by an arrow head. (F-H) Filtration slit proteins podocin (red) and nephrin (green) are expressed at the podocyte cell membranes. (I-K) The podocyte specific transcription factor Wilms’ Tumor-1 (WT1, green) is expressed in podocytes’ nuclei (podocin, red. Nuclei, bleu). (L-N) Phospholipase A2 receptor (PLA2R, red) expression at the podocyte cell membranes (nephrin, green). (O-P’) This cluster was subsetted and merged with the subsetted podocytes from the organoids. Super resolution filtration slit imaging of nephrin in (O-O’) organoid-podocytes and in (P-P’) a morphologically healthy appearing glomerulus of a Munich Wistar Frömter rat.

Podocytes that mirror *in vivo* characteristics as close as possible are required to study NS *in vitro*. Hence, the podocytes in the kidney organoids were thoroughly characterized. The presence of podocyte cell bodies and foot processes as well as a slit diaphragm could be observed at the ultrastructural level (Figure 1D-E). Podocyte specific proteins such as podocin, nephrin, Wilms’ tumor-1 (WT-1), and synaptopodin were are expressed as well (Figure 1F-K, Figure S2). Phospholipase A2 receptor (PLA2R) was also detected in organoid podocytes (Figure 1L-N). PLA2R is the main target receptor of auto antibodies present in the majority of NS patients with primary membranous nephropathy (van de Logt et al., 2019). Slit diaphragms that showed nephrin expression could be detected between podocyte foot processes in organoids, by using super resolution imaging, whereas the morphology closely resembles *in vivo* adult slit diaphragms (Figure 1O-P).

Analysis of single-cell RNA sequencing data revealed three podocyte subclusters, each expressing a distinct profile of podocyte markers, independent of cell cycle phase (Figure 2A-B and Figure S3A). Notably, cluster 3 showed expression of the collagen 4 alpha 3, 4 and 5 (*COL4A3, COL4A4*, and *COL4A5*) genes (Figure 2B). The developmental expression of aforementioned collagen chains is initiated at the capillary loop stage, whereas expression is exclusive for mature podocytes (Abrahamson et al., 2009). The COL4A3 mRNA expression was confirmed in organoid podocytes (NPHS1+) using *in situ* hybridization (Figure 2C). During nephrogenesis, at the s-shaped body phase, podocytes start to secrete vascular endothelial growth factor A (VEGFA), which is essential for podocyte differentiation and attraction of endothelial cell precursors, as a first step towards the formation of a glomerular filtration barrier formation (Abrahamson, 2012; Eremina et al., 2003). All organoid podocyte subclusters expressed VEGFA of which cluster 3 showed the most prominent VEGFA expression (Figure 2D). In fact, VEGFA expression in cluster 3 is 1.5 to 2-fold higher than the expression observed in fetal or adult control podocytes (Figure S3B). The interaction between podocytes and endothelial cells dictates glomerular filtration barrier formation and is essential for glomerular maturation (Schell et al., 2014; Sison et al., 2010), which, however, was not observed in our kidney organoids. Cluster of differentiation 31(CD31), vascular endothelial (VE)-cadherin, plasmalemmal vesicle associated protein-1 (PV-1) and Eps15 Homology Domain-containing 3 (EHD3) where expressed by endothelial capillary structures (Figure 2C, E-G, Figure S4), including luminal spaces. However, the vasculature was interspaced between podocytes, rather than invading as normally occurs at the capillary loop stage during glomerulogenesis *in vivo* (Figure 2E-G).

**Figure 2.**
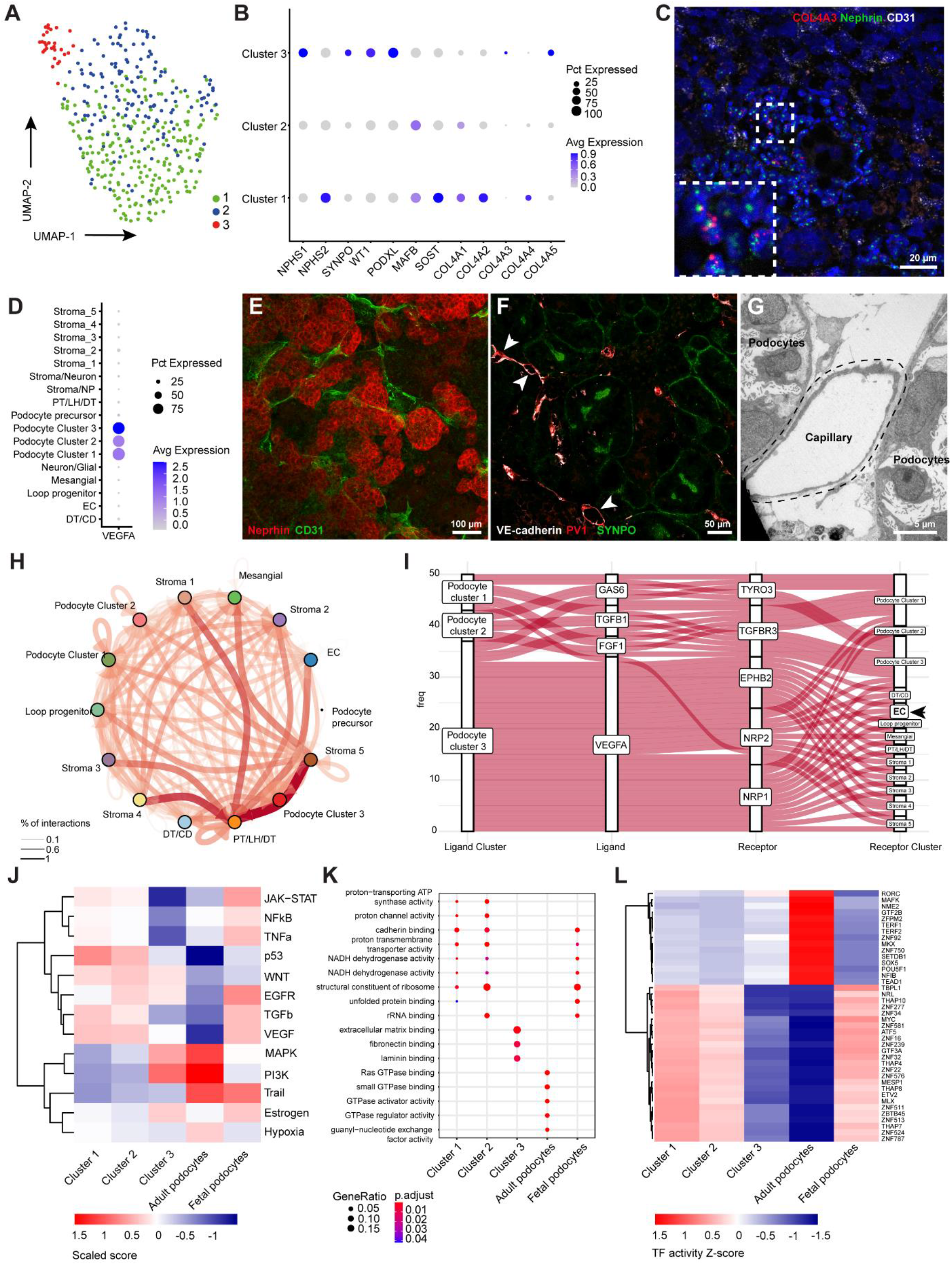
Kidney organoid-derived podocyte subcluster three resembles in part adult podocytes. (A) UMAP integration of 401 podocytes identified three subclusters based on (B) differentially expressed genes. (C) RNAscope analysis showing nephrin (NPHS1), collagen IV alpha three (COL4A3), and endothelial marker cluster of differentiation 31 (CD31) expression. (D) Vascular endothelial growth factor A (VEGFA) expression among the podocyte subclusters. (E) CD31, Vascular endothelial (VE)-cadherin and plasmalemmal vesicle associated protein-1 (PV-1) stained endothelial capillaries are interspaced between (E) NPHS1 and (F) synaptopodin (SYNPO) expressing podocytes. (G) Ultrastructural analysis showing the luminal space of a capillary adjacent to podocytes. (H) Cellcell interactions between all cells present in kidney organoids using CellPhoneDB and CrossTalkeR packages (I) Autocrine and paracrine podocyte ligand-receptor interactions showing a dominant role for VEGFA signaling originating from cluster 3 to VEGF-related receptors in, amongst others, endothelial cell precursors (EC). Analysis was done using CellPhoneDB and CrossTalkeR packages. (J-L) Comparative (J) PROGENy, (K) Gene ontology (GO) terms and (L) DoRothEA analysis showing (J) pathway activity, (K) biological processes and (L) transcription factor activity between the 3 podocyte subclusters, adult and fetal podocytes (adult and fetal podocytes seq data extracted from GEO; GSE118184 and GSE112570, (Laboratory)).

To further dissect cell-cell communication between podocytes and endothelial cells that may aid in explaining observed maturation differences within 3 clusters of podocytes, we performed ligand-receptor analysis using CellPhoneDB and CrossTalkeR packages (Figure 2H) (Efremova et al., 2020; Nagai et al., 2021). Podocyte cluster 3 showed enriched VEGFA interactions with endothelial cell precursors receptors ephrin type-B receptor 2 (EPHB2), Neuropilin-1 and −2 (NRP1 and −2), which could not be identified in the other podocyte clusters (Figure 2I). Concomitantly, using PROGENy (Schubert et al., 2018), podocyte cluster 3 showed an increased PI3K pathway activity (Figure 2J) which, in concert with VEGF signaling, is associated with COL4A3 gene regulation (Chen et al., 2004; Veron et al., 2012). Podocyte cluster 3 mimics adult podocytes most closely when compared to fetal and adult pathway activities that were extracted from the Kidney Interactive Transcriptome database (Figure 2J) (Laboratory). Gene ontology (GO) terms showed distinct biological processes related to extracellular matrix binding in podocyte cluster 3, which were not significantly enriched in the other podocyte clusters or in adult podocytes (Figure 2K). GO terms of podocyte cluster 1 and 2 showed overlap with biological processes active in fetal podocytes (Figure 2K). To infer transcription factor (TF) activity differences between the podocytes clusters that may aid in resolving molecular cues involved in podocyte maturation, we used the DoRothEA computational package (Garcia-Alonso et al., 2019). We found that podocyte cluster 1 and 2 resemble a fetal podocyte TF profile whereas the transcriptional circuit in podocyte cluster 3 showed partial overlap with adult podocytes (Figure 2L). However, some emerging TFs, like TEA domain transcription factor 1 (TEAD1) and nuclear factor I B (NFIB), involved in cellular differentiation and tissue homeostasis are less active in podocyte cluster 3 compared to adult podocytes (Figure 2L) (Adam et al., 2020; Rinschen et al., 2017; Sabo et al., 2014). Of note, zinc finger protein 750 (ZNF750) was less enriched in cluster 3 podocytes compared to adult podocytes. Normally, ZNF750 represses kruppel like factor 4 (KLF4), a known epigenetic modulator involved in podocyte differentiation (Hayashi et al., 2014). Also, SRY-Box transcription factor 5 (SOX5) known to co-activate WT1 (Dong et al., 2015), shows a limited activity in cluster 3. Altogether, we show that essential podocytes markers are expressed in the entire kidney organoid podocyte population, whereas we identified a subpopulation of podocytes that resemble adult podocytes at the transcriptional level.

Next, we compared the performance of the current organoid model compared with a frequently used human 2D *in vitro* podocyte cell line, i.e. a widely used conditionally immortalized podocytes cell line (ciPOD) (Saleem et al., 2002)). Morphologically, ciPODs do not show the typical appearance of a “floating” cell body with primary and secondary (foot process) structures (Figure 3B). Notably, within the organoids, podocytes highly resembled *in vivo* podocytes (Figure 1E’ and 3A-C). Transcriptome analysis revealed that organoid podocytes much more resemble *in vivo* podocytes than ciPODs (Figure 3D), in particular when evaluating typical podocyte markers such as NPHS1, NPHS2, WT1, PODXL, and SYNPO. Altogether, the organoid podocytes in part resemble the transcriptome profile of human adult kidney podocytes, which includes the expression of typical podocyte markers being essential to study NS pathology *in vitro*.

**Figure 3.**
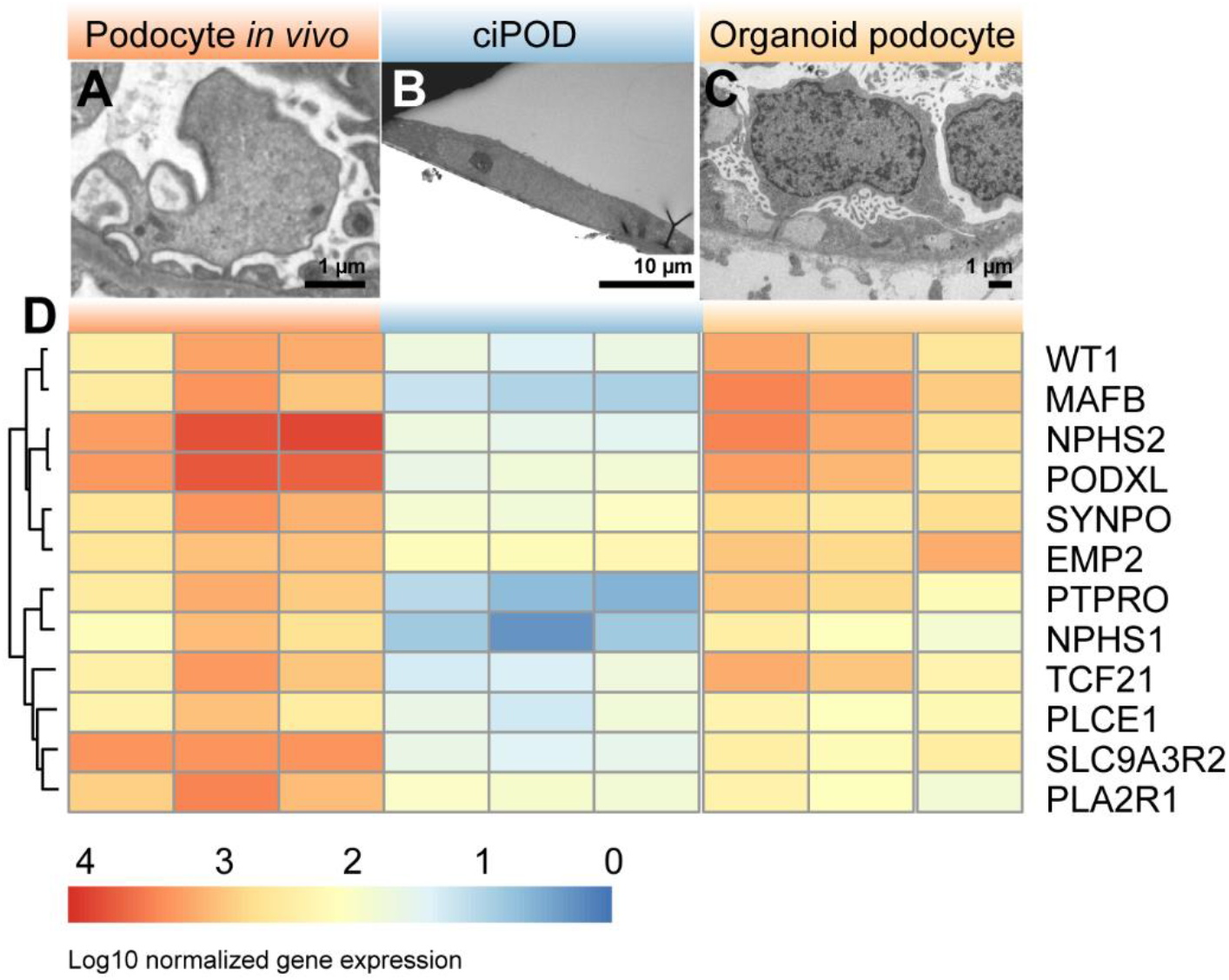
Organoid podocytes show superior morphology and transcriptome as compared to 2D podocytes *in vitro*. (A) Ultrastructural analysis of normal human podocytes *in situ*, (B) conditionally immortalized podocytes (ciPOD) and (C) organoid podocytes. (D) Heatmap of bulk RNA sequencing data showing the expression signature of podocyte specific markers in normal human glomeruli, ciPOD and organoid podocytes.

### Kidney organoids of a patient with a NPHS2-mutant display impaired protein processing towards the cell membrane that was abrogated after genetic correction

Podocin is known to play an important role in slit diaphragm organization. Intact podocin is required for nephrin membrane trafficking, ultimately leading to oligomerization of podocin clusters and nephrin to develop filtration slits. NS patients with mutated *NPHS2* are characterized by an aberrant nephrin expression, thereby emphasizing the crucial role of podocin in podocyte physiology (Jaffer et al., 2011; Zhang et al., 2004). Here, we used kidney organoids to model congenital nephrotic syndrome of a *NPHS2*-mutant.

A patient with congenital nephrotic syndrome was diagnosed with the compound heterozygous mutations p.Arg138Gln (exon 3) and p.Asp160Tyr (exon 4) in the podocin (*NPHS2*) gene. Erythroblasts from this patient were successfully reprogrammed into iPSC and cultured into organoids. Podocin protein expression was nearly absent on the cell membrane and although nephrin expression was present, its distribution was aberrant (Figure 4A-C), which is in line with observations in NS patients (Zhang et al., 2004). Correction of the exon 3 mutation by CRISPR/Cas9 resulted in restored podocin protein expression colocalizing with nephrin experssion at the cell membrane in organoids (Figure 4D-F). Bulk transcriptomics of organoid NPHS1+ podocytes showed a distinct podocyte mRNA expression profile for both mutant and repaired organoid podocytes, for example SYNPO, MAFB, WT1, NPHS1 and NPHS2 expression (Figure 4G), thereby resembling the transcriptome of isolated human glomeruli (FigureS5A). The NPHS2 gene expression is not affected by the NPHS2 mutations. Both NPHS2 mutations result in protein retention in the endoplasmic reticulum and do not interfere with mRNA expression (Nishibori et al., 2004; Tory et al., 2014).

**Figure 4.**
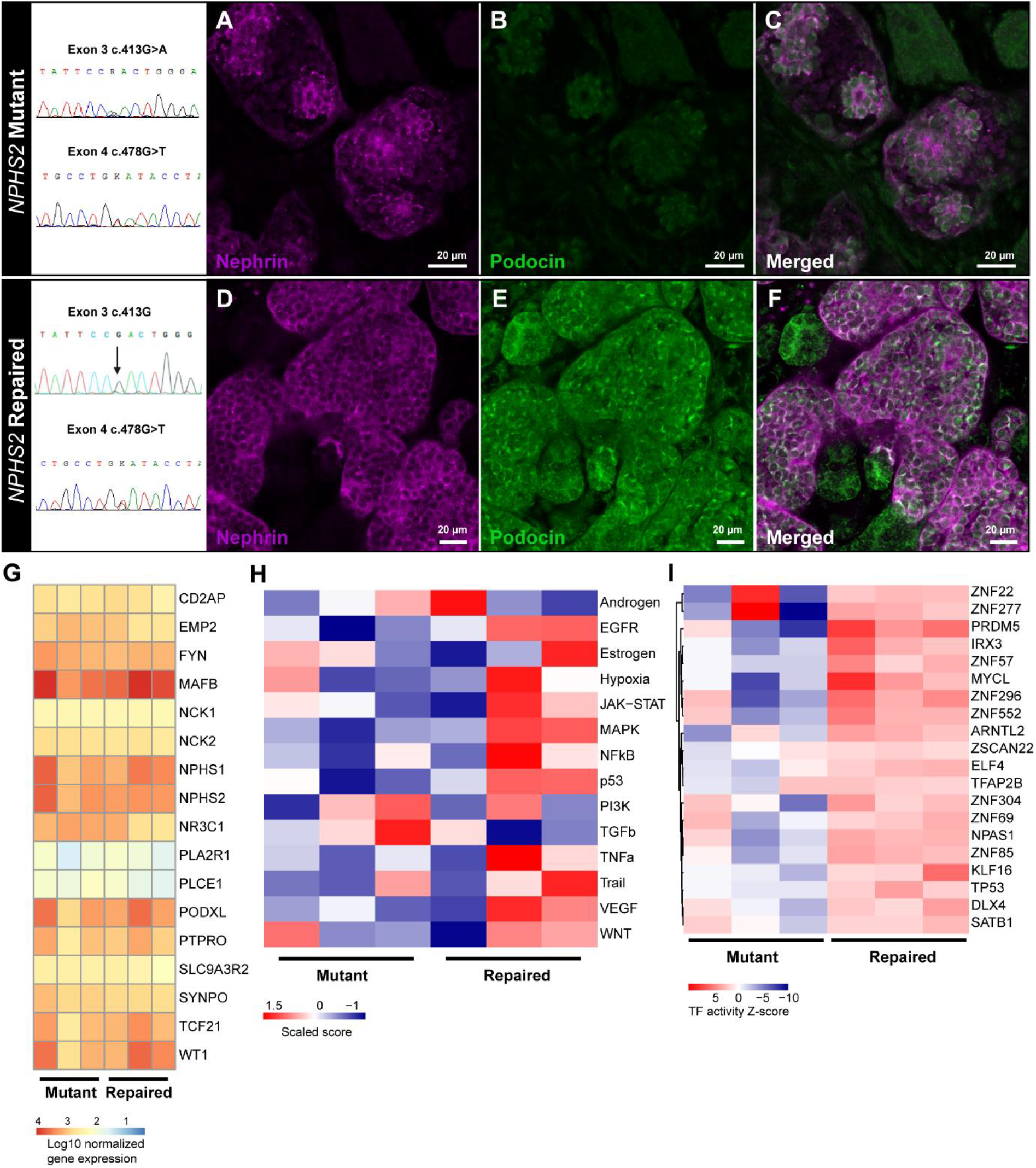
Congenital nephrotic syndrome modeling of patient-derived *NPHS2* mutant and genetically repaired iPSC kidney organoids. (A-F) Nephrin (magenta) and podocin (green) expression in (A-C) *NPHS2* mutant and (D-F) genetically repaired kidney organoids. (G) Comparative bulk transcriptomics of podocyte specific genes in mutant *versus* repaired organoid podocytes. (H) Comparative PROGENy pathway activity analysis and (I) DoRoThea transcription factor activity of mutant *versus* repaired organoid podocytes.

*In vivo*, podocytes are the major VEGF producers in the glomerular compartment, which acts via autocrine and paracrine signaling to maintain the glomerular filtration barrier (Chen et al., 2004). Autocrine VEGF-A signaling regulates slit diaphragm proteins including podocin (Guan et al., 2006). In line with this, we showed by using PROGENy, that repaired organoid podocytes displayed enriched VEGF pathway activity compared to mutant organoid podocytes (Figure 4H and Figure S5B). Repaired organoid podocytes closely resemble VEGF pathway activity levels of fetal podocytes, whereas mutant organoid podocytes differ from fetal podocytes (Figure S5B). We identified that fibroblast growth factor (FGF)-FGF receptor 4 (FGFR4) signaling is enhanced in repaired organoid podocytes compared to mutant organoid podocytes (Figure S5C – S6, (Liberzon et al., 2015)). FGF signaling is likely to be involved in podocyte differentiation and plays a role in podocyte recovery following injury (Reiser et al., 2014). All up- and downregulated genes in mutant *versus* repaired organoids are shown in Figure S5 C-D.

The transcription factor activity of repaired organoid podocytes showed a distinct signature as compared to mutant organoid podocytes (Figure 4I). The top 20 enriched TFs include, amongst others, PR domain zinc finger protein 5 (PRDM5), E74-like ETS transcription factor 4 (ELF4) and zinc finger protein 69 (ZNF69) which are involved in the regulation of genes involved in podocyte physiology (Sharmin et al., 2016; Wu et al., 2018). In addition, zinc finger protein 277 (ZNF277) is enriched in repaired organoid podocytes which is involved in the regulation of synaptopodin as well as CD2AP (Lu et al., 2017). Altogether, we successfully modeled congenital NS in organoids. The repaired *NPHS2* mutation resulted in restored podocin expression and nephrin localization, as well as the rescue of pathway and TF activities essential in podocyte physiology.

### Idiopathic NS modeling using kidney organoids shows downstream slit diaphragm signaling events and granule formation in podocytes

The protamine sulfate model is a known *in vivo* model to specifically induce podocyte injury (Verma et al., 2016). Protamine sulfate abolishes the negative charge on podocytes, thereby inducing foot processes effacement accompanied by downstream slit diaphragm signaling, cytoskeleton rearrangements and activation of injury associated mechanisms. The protamine sulfate model is reversible by heparin treatment, which ultimately restores podocyte charge, the cytoskeleton and foot processes (Verma et al., 2016).

Organoids treated with protamine sulfate showed clear podocyte cytoskeleton rearrangements (Figure 5C, 5Q) and podocyte injury, as shown by vacuole formation and cell degeneration at the ultrastructural level (Figure 5G) as compared to control (Figure 5A-B and 5E-F). The PS-induced injury could largely be reversed by heparin treatment (Figure 5D, 5H, and 5Q). iPSC differentiated into 2D podocytes, from the same iPSC as used for organoids, as well as ciPOD also showed cytoskeleton rearrangements as a result of protamine sulfate exposition (Figure 5K and Figure 5O), which also showed partial recovery after heparin treatment (Figure 5L, 5P). Interestingly, only the organoids showed nephrin and protamine sulfate induced phospho-nephrin expression upon western blotting (Figure 5R, 5T), which is indicative for slit diaphragm signaling following injury, whereas neither in 2D iPSC-derived podocytes nor in ciPODs any nephrin expression could be detected (Figure 5R-S).

**Figure 5.**
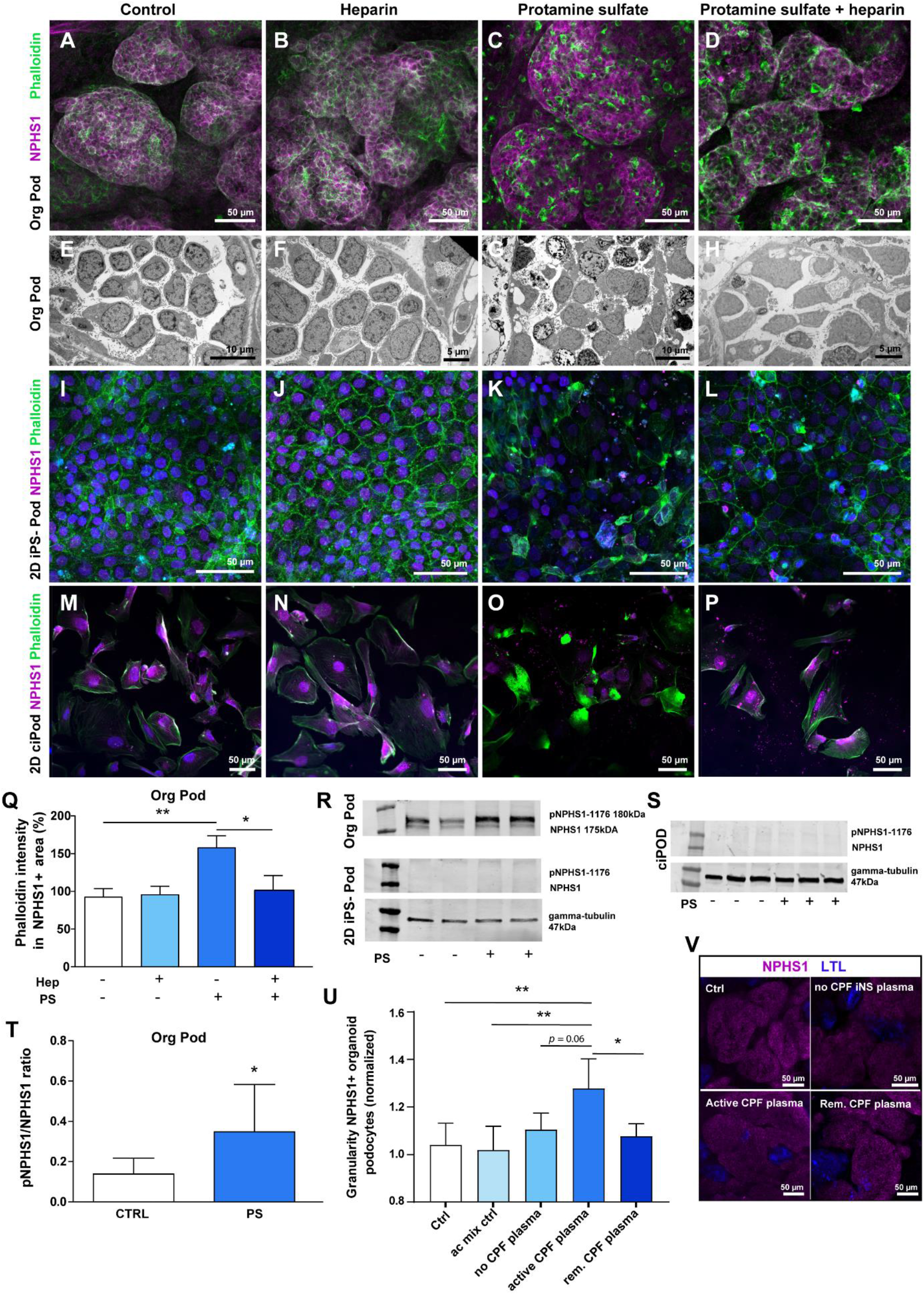
Kidney organoids are the preferred model of choice to model idiopathic nephrotic syndrome. (A-D) Podocyte cytoskeleton (phalloidin (green), NPHS1 (magenta)) and (E-H) ultrastructure analysis of protamine sulfate treated and heparin rescued kidney organoid podocytes. (I-L) Podocyte cytoskeleton analysis (phalloidin (green), NPHS1 (magenta)) of protamine sulfate treated and heparin rescued 2D iPSC-derived podocytes and (M-P) ciPODs. (Q) Quantification phalloidin intensity in protamine sulfate treated and heparin rescued kidney organoid podocytes. *=p<0.05, ** = p<0.01. (R-S) pNPHS1 and NPHS1 protein expression analysis of protamine sulfate treated kidney organoids, 2D iPSC-derived podocytes and (S) ciPODs. Gamma tubulin was used as reference protein. (T) Western blotting semi-quantification pNPHS1/NPHS1 ratio of protamine sulfate treated kidney organoids. *=p<0.05. (U) Cytoplasmic granularity analysis in NPHS1-sorted organoid podocytes, isolated from kidney organoids treated with 10% (v/v) iNS plasmapheresis plasma (circulating permeability factor (CPF) free, active CPF and remission CPF plasma), control (E6 medium) and anti-coagulation mix (ac mix ctrl) for 4h (N=2 independent experiments, triplicate per condition). *=p<0.05, **=p<0.01. (V) NPHS1 (magenta) and LTL (blue) expression following plasma treatments in kidney organoids.

Next, as a proof of concept, we showed that organoid podocytes respond to CPFs present in plasma from patients with NS recurrence after kidney transplantation (Figure 5U). We treated organoids for 4h with plasma collected from plasmapheresis during the active course of the disease and plasma obtained after remission. Control plasmapheresis plasma was collected from an iNS patient after kidney transplantation, with no disease recurrence (plasma presumably free of CPFs). By flow cytometry analysis, we could show that NPHS1+ organoid podocytes exhibited increased cytoplasmic granules formation when exposed to active plasma as compared to control and remission plasma (Figure 5V). Podocyte granule formation is an acknowledged marker for podocyte injury (den Braanker et al., 2021; Seckin et al., 2017; Yamamoto-Nonaka et al., 2016). Podocyte morphology and nephrin protein expression were not affected after plasma exposure (Figure 5V and Figure S7). Our data indicate that organoid podocytes could be a useful model to investigate podocyte pathophysiology activated by CPFs and, ultimately, may contribute to identify CPFs involved in iNS.

### Intermodel evaluation affirms the need for 3D organoids to accurately model NS

To further investigate the preferred model of choice, i.e. 2D iPSC-derived podocytes *versus* organoids, to study NS, the podocin mutant, and repaired iPSC lines were used evaluated in both models. Mutant and repaired organoids were exposed to protamine sulfate and/or treated with heparin. In line with our previous results, these organoids showed cytoskeleton rearrangements as a result of protamine sulfate exposure (Figure 6A, E), which was only significantly reversed upon heparin treatment in podocin repaired organoids (Figure 6E).

**Figure 6.**
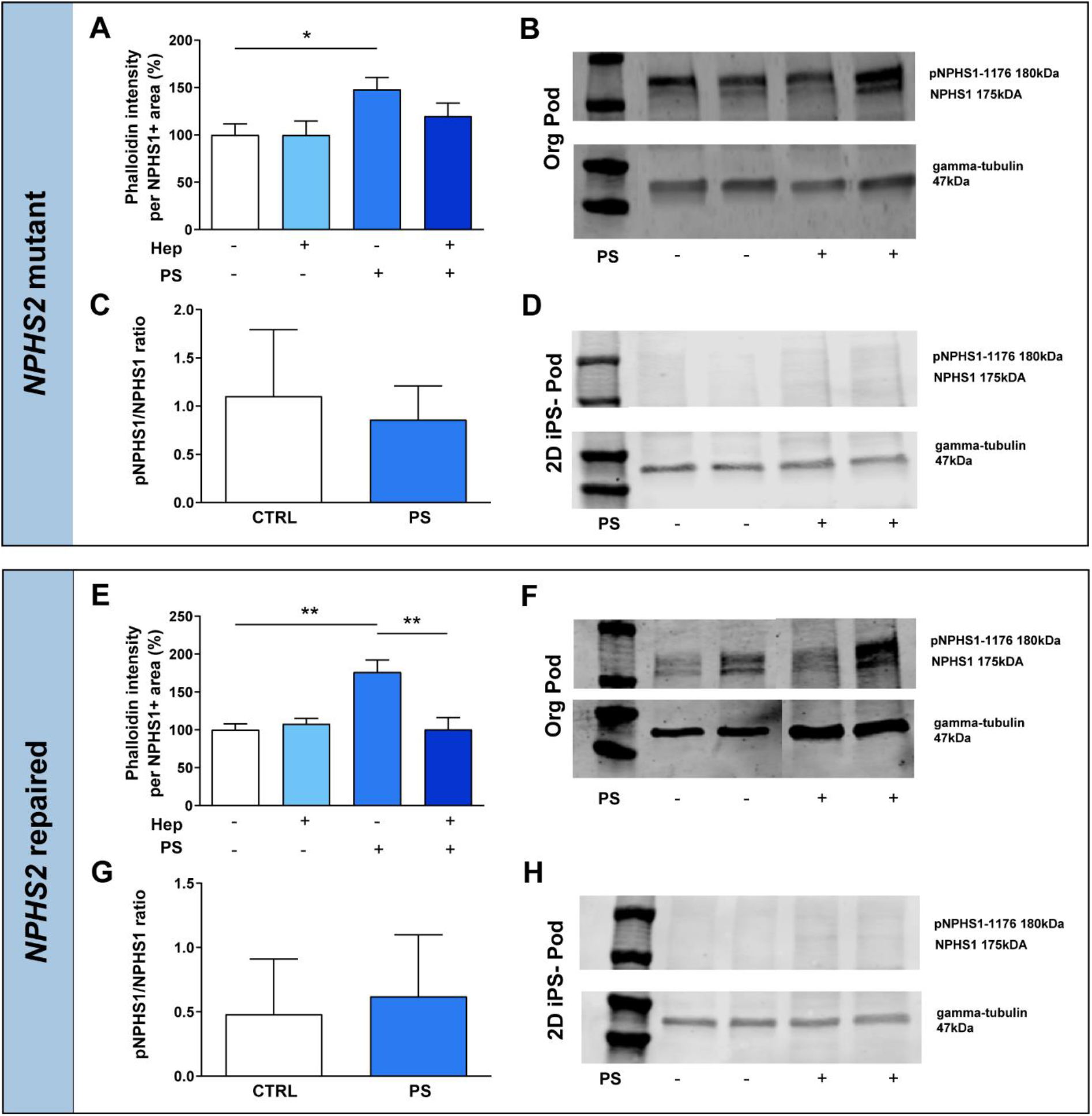
Intermodel evaluation supports the need for kidney organoids to model idiopathic nephrotic syndrome. Quantification of phalloidin intensity in protamine sulfate treated and heparin rescued podocytes in (A) mutant and (E) repaired kidney organoids. *=p<0.05, ** = p<0.01. pNPHS1 and NPHS1 protein expression analysis of protamine sulfate treated (B) mutant and (F) repaired kidney organoids. Western blotting quantification pNPHS1/NPHS1 ratio of protamine sulfate treated (C) mutant and (G) repaired kidney organoids. pNPHS1 and NPHS1 protein expression analysis of protamine sulfate treated 2D (D) mutant and (H) repaired iPSC-derived podocytes. Gamma tubulin was used as reference protein.

Moreover, pNPHS1 expression in both organoid lines was not significantly induced upon protamine sulfate treatment. Nevertheless, a slight trend in pNPHS1 expression in the repaired line could be observed (Figure 6B-C and Figure 6F-G). iPSC, both mutant and repaired, differentiated in 2D towards podocytes responded to protamine sulfate resulting in injury and recovered upon heparin treatment according to actin cytoskeleton expression (Figure S8). However, no nephrin and pNPHS1 expression was observed in 2D iPSC-derived podocytes (Figure 6D, H), which complies with our earlier results. These results support that 3D organoids are the preferred model of choice to accurately study podocytes in culture.

## Discussion

In the present study, we successfully modeled congenital (*NPHS2* mutant) and idiopathic NS using kidney organoids. Using iPSCs derived from a patient with two heterozygous podocin mutations, we could show the absence of podocin expression and aberrant nephrin localization, which was rescued by CRISPR/Cas9 gene editing in kidney organoids. As a result, the VEGFA pathway and transcription factors, essential in podocyte physiology, were restored. Idiopathic NS was modeled in organoids using plasma from a patient with post-transplant disease recurrence which resulted in podocyte injury. Moreover, we induced iNS-associated podocyte injury using protamine sulfate. We report that kidney organoids show superior morphology, expression of podocyte markers and activate downstream signaling cascades upon injury which are absent in 2D iPSC-derived podocytes and human ciPOD, thereby making kidney organoids the preferred model to study NS *in vitro*.

NS pathophysiology is complex and the molecular mechanisms underlying the disease are still poorly understood, mainly due to a lack of accurate human models. From previous studies it is known that the podocyte is the main culprit in NS, consequently affecting GBM integrity and leading to endothelial cell injury (Hansen et al., 2020; Rinschen et al., 2018; Smeets et al., 2011; Wharram et al., 2005). For decades, murine and human 2D conditionally immortalized podocytes (ciPOD) have been used to unravel NS pathophysiology. Although these models have expanded our knowledge regarding podocytopathies in NS, we and others showed that *in vitro* 2D conditionally immortalized podocyte models do not meet *in vivo* podocyte characteristics (Hagmann and Brinkkoetter, 2018; Hale et al., 2018; Rinschen et al., 2018; van den Broek et al., 2021). Here, we showed that kidney organoids are a promising tool to study both congenital and idiopathic nephrotic syndrome since organoid podocytes express essential proteins, with a morphology closely resembling *in vivo* podocytes, and respond similarly to injury triggers. The 3D organoids show a superior expression profile compared to the 2D iPSC-derived podocytes. In 2D, nephrin is absent or the localization is still intracellular rather than between the interdigitating foot processes. Also, downstream filtration slit signaling is absent in injured 2D iPSC-derived podocytes, emphasizing the limited differentiation properties of podocytes in 2D culture. It is known that 3D glomeruli-like structures isolated from organoids maintain marker expression for 96h, whilst 2D podocytes migrating from these organoid glomeruli show a clear alteration in the their transcriptome (Hale et al., 2018). There is an unmet need for better in vitro models to recapitulate the in vivo glomerular filtration barrier, with emphasis on the podocyte. The 3D configurations and the role of mechanical as well as extracellular matrix (ECM) cues that influence cellular (de)differentiation to develop accurate organoid models need to be addressed in order to establish improved glomerular models *in vitro*. The effect of ECM cues and material stiffness on organoid differentiation was nicely shown by Garetta et al and also by Geuens and co-workers (Garreta et al., 2019; Geuens et al., 2021). Organoids encapsulated in soft biomaterials showed accelerated differentiation, less fibrosis upon aging, and reduced off-target cell populations, all aspects that will aid in signaling to enhance the maturation of podocytes but also other kidney segments like the proximal tubule and the collecting duct.

Kidney organoids hold great promise as the next step in NS modeling, nevertheless, tissue maturity and the presence of off-target cells are often being criticized. To overcome maturation challenges, organoid transplantation *in vivo* or onto chicken chorioallantoic membranes can be performed (Garreta et al., 2019; Sharmin et al., 2016; van den Berg et al., 2020; van den Berg et al., 2018). Organoid maturation can be achieved by the use of renal subcapsular transplantation of human kidney organoids in immunodeficient mice for the adaptive immune system, which results is in size-selective filtration across organoid glomerular filtration barrier (van den Berg et al., 2020). This transplantation approach emphasizes the potential of organoids to develop a mature glomerulus to study a wide range of glomerular disorders, though animals would still be required which hampers high-throughput screens and is in conflict with the current attention for animal welfare. Altogether, when focusing on podocytopathies like NS, the current organoids we present in this study represent a useful tool to model idiopathic and congenital injury responses.

The human adult podocyte signature is not yet accomplished in kidney organoids. Computational analysis revealed heterogeneity throughout the organoid-podocyte populations based on collagen IV alpha chain expression. It is known that at the comma, S-shaped, and early capillary loop stage of glomerulogenesis, the podocyte progenitors express collagen IV alpha chains 1 and 2. This collagen IV network is replaced by collagen IV alpha chains 3, 4, and 5 at the beginning of capillary loop stages and ultimately these chains are solely expressed by mature podocytes (Abrahamson et al., 2009). The exact mechanism of the collagen IV isoform switch remains elusive, though it has been described to aid in podocyte differentiation and is likely under the control of VEGF, which in turn is regulated by, amongst others, PI3K signaling (Chen et al., 2004; Sison et al., 2010; Veron et al., 2012). In line with this, we showed that podocyte subcluster three had increased collagen IV alpha 3 and VEGF expression as well as PI3K signaling compared to the other two podocyte subclusters. Moreover, solely cluster three shows ligand-receptor interactions with endothelial cell precursor receptors, as a potential first step towards glomerular filtration barrier formation, thereby confirming that cluster three is more mature compared to the other two podocyte clusters. Cluster 3 resemble in part adult podocytes in terms of collagen IV alpha 3 expression. The data we collected in this study could aid in resolving what culture conditions we would need to get a fully mature podocyte population in organoids.

The PI3K signaling pathway, which regulates VEGF signaling, can be induced by tyrosine kinases like epidermal growth factor receptor (EGFR) (Pages and Pouyssegur, 2005), which is achieved in our kidney organoid culture protocol by EGF supplementation. Despite the EGFR stimulation, the consequent PI3K activation did not result in homogenous VEGF expression levels as VEGF expression varies between the podocyte clusters we identified in our organoids. Also the TF signature in podocyte cluster 3 showed closest, but still incomplete, resemblance with adult podocytes. To accelerate differentiation of podocytes, VEGF signaling is likely one of the major actors (Eremina et al., 2003). When focusing on VEGF signaling, a magnitude of factors are involved to regulate VEGF signaling. The promoter region of VEGF contains multiple trans-activating factors and responsive elements (Pages and Pouyssegur, 2005), like hypoxia inducible factor-1 (HIF1), that might open avenues to transcriptionally stimulate VEGF expression throughout all podocytes present in the organoids. Notably, organoids are cultured under normoxic conditions, which differs from glomerulogenesis *in vivo* which occurs under hypoxic conditions (Ryan et al., 2021). A hypoxic environment will activate HIF transcription factors, regulate VEGF signaling, and consequently stimulate podocyte differentiation (Ryan et al., 2021). Mimicking the delicate balance of VEGF signaling during the different stages of nephrogenesis will be challenging to accurately tune *in vitro* (Eremina et al., 2003). Obviously, podocyte maturation not only depends on VEGF signaling but relies on a plethora of factors. For instance, podocyte maturation also relies on the concerted actions with other cell types, such as podocyte-macrophage crosstalk (Munro et al., 2019), fluid flow including nutrient exchange as well as microenvironmental ECM cues (Garreta et al., 2019; Geuens et al., 2021; Homan et al., 2019; Ryan et al., 2021; van den Berg et al., 2020).

In conclusion, we report successful patient-based idiopathic and congenital nephrotic syndrome modeling in iPSC-derived human kidney organoids. We showed that 3D kidney organoids, and in particular a specific podocyte cluster in part resemble adult podocytes, thereby being the preferred model of choice in NS modeling, as confirmed in three iPSC lines, compared to 2D cultured podocytes. Kidney organoids represent a powerful model to study NS disease mechanisms, have the potential to identify pathogenic CPFs in patients’ plasma, and allow for the study of functional genomics. Altogether, kidney organoids will aid in resolving podocytopathies and may be used for developing novel effective therapies to treat NS. Once a mature glomerular filtration barrier is developed *in vitro*, this high-end platform could be used beyond studying podocytopathies and may have the potential to become an accurate model in resolving the pathophysiology of a wide range of glomerular disorders.

## Author Contributions

J.J., M.F.S. and B.S. designed the study. J.J., B.T.B., M.B. and K.C.R cultured the iPSC-derived kidney organoids. J.J., B.T.B., M.B., GDG, F.M. Q.N., R.W and H.M. performed tissue processing, IF stainings, Western Blotting as well as RNAscope experiments. B.W. carried out electron microscopy analysis. E.S., T.N., R.J.M., J.F.M.W., J.V. contributed tissue specimens and plasma material. C.K. performed data acquisition experiments for single cell sequencing. C.K. and R.K. supported singe cell sequencing. M.B. carried out the single cell sequencing data analyses, and was assisted by J.S.N. and I.G.C.. N.E. and V.D. performed super resolution imaging. J.J., B.T.B. and M.B. arranged the Figures. J.J., B.T.B., M.B, M.F.S. and B.S. wrote the manuscript. J.F.M.W., N.K., R.J.M., T.N., J.V. K.C.R, C.K. and R.K. edited the manuscript and advised on data analysis and interpretation. All authors read and approved the final manuscript.

## Acknowledgments

J.J. is supported by the Netherlands Organization for Scientific Research (NWO Veni grant no: 091 501 61 81 01 36) and the Dutch Kidney Foundation (grant no 19OK005). B.T.B is supported by grants awarded to R.J.M.: the Netherlands Organization for Scientific Research (NWO Off road grant no: 451001034) and the Dutch Kidney Foundation (grant No: 18OKG05 and 18OI14). M.B. is supported by the Dutch Kidney Foundation (grant no 19OK005). M.F.S. is supported by the Dutch Kidney foundation (grant 15OKG16) and The Netherlands Organization for Scientific Research (NWO VIDI grant: 016.156.454). B.S. is supported by the Dutch Kidney foundation (grant 14A3D104) and The Netherlands Organization for Scientific Research (NWO VIDI grant: 016.156.363). J.S.N. and I.G.C. are supported by the E:MED Consortia Fibromap funded by the German Ministry of Education and Science (BMBF).

We would like to thank the Stem cell technology center, Radboudumc, Nijmegen, the Netherlands and the iPS Core Facility, Erasmus Medical Center, Rotterdam, the Netherlands, for generating iPSC and performing the gene editing. We would like to thank the Radboudumc Technology Centers bioinformatics, microscopy and flow cytometry for their support.

## Materials and Methods

### Ethical statement

Human adult skin fibroblasts derived from a healthy volunteer and erythroblasts from a pediatric patient suffering from congenital nephrotic syndrome, after giving informed consent, were used to generate iPSC. This study was conducted in accordance with the Helsinki Declaration as revised in 2013. Permission for the creation and use of iPSCs in this study was obtained from the local ethical commission for human-related research of the Radboud university medical center, Nijmegen (approval numbers: 2015-1543 and 2006-048). Human kidney tissue was used for Immunostainings and EM analysis. Ethical permission for the use of archived human kidney material was given by the local commission for human-related research of the Radboud university medical center, the Netherlands (approval number: 2018-4086).

### Patient and healthy control plasma

This study was approved by the regional medical-ethical committee (Arnhem-Nijmegen), under file number 09/073, and has been carried out in compliance with the Helsinki declaration as revised in 2013. All human participants signed informed consent prior to the study, as specified in the *ICMJE* Recommendations.

### Reagents and resources

In Supplemental table S5 reagents and resources are listed (e.g. antibodies, chemicals, commercial assays, software and algorithms).

### Cell culture

#### iPSC generation

Healthy control iPSC were generated using fibroblasts from a healthy volunteer at the Radboudumc Stem Cell Technology Center through the use of lentiviral vectors, previously described by Warlich et al. (Warlich et al., 2011). In short, *E. coli* were used to clone lentiviral vectors containing the transcription factors Oct4, Klf4, Sox2 and cMyc together with the fluorescent marker dTomato (pRRL.PPT.SF.hOKSMco-idTom.pre.FRT). HEK293T cells were then used for viral production using packaging (pCMV-VSV-G) and envelope (pCMV-dR8.91.) vectors, respectively. Fibroblasts were transduced with viral supernatants and reseeded on plates with inactivated mouse embryonic fibroblast (MEF) feeder cells to support the growth and pluripotency of iPSCs. Emerging iPSC colonies were picked, expanded and assessed for activation of stem cell markers to confirm pluripotency.

Mutant iPSC, containing 2 mutations in the podocin encoding gene (NPHS2; Chr1(GRCh37); NM_014625.3:c.413G>A (p.(Arg138Gln)); heterozygote and c.478G>T (p.(Asp160Tyr); heterozygote) were generated from erythroblasts from a pediatric patient using the CytoTune™-iPS 2.0 Sendai Reprogramming Kit (ThermoFisher Breda, The Netherlands) according to the manufacturers protocol. In short, erythroblasts were transduced using the CytoTune™-iPS 2.0 Sendai Reprogramming Kit and reseeded on plates with inactivated mouse embryonic fibroblasts to support the growth and pluripotency of iPSCs. Emerging iPSC colonies were picked, expanded and assessed for activation of stem cell markers to confirm pluripotency.

#### iPSC maintenance culture

iPSCs were cultured using Essential™ 8 (E8) medium (Gibco, ThermoFisher) supplemented with E8 supplement (50X, Gibco) and 0.5% (v/v) antibiotic-antimycotic (Gibco) on Geltrex-coated (ThermoFisher) cell culture plates (Greiner, Alphen aan den Rijn, the Netherlands). Upon 70-90% confluency iPSCs were washed 3 times with PBS and subsequently passaged in colonies using 0.5 mM EDTA (ThermoFisher) in PBS for 5 minutes at room temperature. For cell seeding, iPSCs were washed 3 times with PBS and subsequently disassociated into single cells using TrypLE Select Enzyme (ThermoFisher) for 2 minutes at 37°C.

#### ciPOD culture

The conditionally immortalized podocyte cell line (ciPOD) used in this study was kindly provided by Prof. Saleem (Bristol, UK)(Saleem et al., 2002) ciPODs were cultured using uncoated 75 cm^2^ culture flasks (Greiner) at 33 °C with 5% (v/v) CO_2_ in DMEM/F12 (Gibco) supplemented with 5 μg/mL insulin, 5 μg/mL transferrin, 5 ng/mL selenium (Sigma-Aldrich, Zwijndrecht, the Netherlands), 10% heat inactivated FCS (Gibco) and 1% (v/v) penicillin/streptomycin (Gibco). Passaging and seeding of cells was performed by washing cells with PBS and subsequently detaching them using Accutase® (Sigma-Aldrich), according to the manufacturer’s protocol. ciPODs were seeded at 62,500/cm^2^ in black flat bottom 96-wells plates (Corning, Corning, New York) for immunofluorescent imaging or 6-wells plates (Greiner) for protein and RNA isolation and were incubated at 37°C with 5% (v/v) CO_2_, which initiated differentiation of the ciPODs.

### Differentiation protocols

#### 2D podocyte differentiation

An adapted protocol based on Rauch *et al*. and Musah *et al*. (Musah et al., 2017; Rauch et al., 2018) was used to differentiate iPSC into podocytes. iPSCs were seeded at 15,000 cells/cm^2^ on Geltrex-coated cell culture plates in E8 medium supplemented with E8 supplement, 0.5% (v/v) antibiotic-antimycotic and 1X RevitaCell (Gibco). One day later, defined as day 1, differentiation of iPSCs into metanephric mesenchyme was started by adding Essential™ 6 (E6) medium (Gibco), supplemented with 15 ng/mL human recombinant bone morphogenetic protein 7 (RD system, Minneapolis, Minnesota), 10 ng/mL human recombinant activin A (RD systems), 100 nM retinoic acid (Sigma-Aldrich) and 1X non-essential amino acids (Gibco). At day 11, podocyte differentiation was started by culturing cells in E6 medium supplemented with 1X non-essential amino acids and 10 ng/mL vascular endothelial growth factor subtype A (RD systems). At day 20, iPSC-derived podocytes were mature and used for experiments.

#### 3D organoid differentiation

iPSCs were seeded on Geltrex-coated cell culture plates using E8 medium supplemented with E8 supplement, 0.5% (v/v) antibiotic-antimycotic and 1X RevitaCell at a density between 18,750 - 22,900 cells/cm^2^. Twenty-four hours after seeding, culture medium was replaced by E6 medium supplemented with 6 μM CHIR (Tocris, Bio-Techne, Abingdon, United Kingdom) to initiate differentiation (defined as day 0). From day 3 onwards, half of the wells were cultured in E6 medium supplemented with 200 ng/mL FGF9 (RD systems) and 1 μg/ml heparin (Sigma-Aldrich) until day 7 (anterior intermediate mesoderm (AM)), whereas the other wells remained on E6 medium with CHIR until day 5 (posterior intermediate mesoderm (PM)). From day 5 onward, all cells were cultured in E6 medium with FGF9 and heparin. At day 7, cell were harvested using trypsin-EDTA and a cell suspension was made consisting of AM and PM cells in a 1:2 ratio. Cells aggregates, consisting of 300K cells, were transferred to Transwell™ plates (Corning) and incubated for at least 1 hour at 37°C, 5% (v/v) CO_2_, with E6 medium supplemented with 5 μM CHIR to induce self-organizing nephrogenesis. After 1 hour, medium was replaced for E6 medium supplemented with FGF9 and heparin. At day 7+5 (i.e. 5 days later), medium was changed to E6 supplemented with EGF (10 ng/ml, RD systems) and BMP7 (50 ng/ml, RD systems). Organoids were used for experiments at day 7+17 or 7+18.

### Protamine sulfate exposure and heparin rescue

Organoids, ciPODs and 2D iPSC-derived podocytes were exposed to protamine sulfate treatment and rescued with heparin. Briefly, protamine sulfate (2 mg/ml, Sigma-Aldrich) was dissolved in E6 medium and organoids and 2D cell cultures were exposed at 37°C, 5% (v/v) CO_2_ for 2 hours. After 2 hours, cells (organoids and 2D cells) exposed to protamine sulfate were processed for protein isolation and immunofluorescent staining, as described in a following sections. Sequentially, protamine sulfate injured organoids and 2D cell cultures were exposed to heparin (800 μg/ml in E6, heparin sodium salt from porcine intestinal mucosa, Sigma) at 37°C, 5% (v/v) CO_2_ for 2 hours. Ultimately after 2 hours, these organoids and cell cultures (groups: control, heparin control, protamine sulfate plus heparin treatment) were also processed for protein isolation and immunofluorescent staining.

### Preparing paraffine sections of human iPSC-derived kidney organoids

iPSC-derived kidney organoids were cut from the Transwell™ filter and fixated in 4% (v/v) formalin on ice for 20 minutes. Fixed iPSC-derived kidney organoids were stripped of the filter membrane using a scalpel. Multiple organoids from similar conditions were stacked on top of each other and embedded in a cryomold (Tissue-Tek, Sakura Finetek Europe B.V., Alphen aan de Rijn, the Netherlands) using 2.25% (w/v) agarose gel (ThermoFisher). After embedding for 5 minutes at 4°C, the iPSC-derived kidney organoids were transferred to embedding cassettes and paraffinized. After paraffinization, iPSC-derived kidney organoids were cut at a thickness of 4 μm using a cryo-microtome and mounted on FLEX IHC Microscope Slides (DAKO, Agilent Technologies, Amstelveen, Netherlands).

### Immunofluorescence staining

#### Paraffine tissue

Using a series of xylol (2x) and 100% (v/v) ethanol (3x), paraffine slides were deparaffinized. Antigen retrieval was performed by boiling slides in tris-buffered EDTA (VWR Chemicals, Amsterdam, the Netherlands) for 10 minutes in a microwave. Primary (1:100) and secondary (1:200) antibodies were diluted in PBS containing 1% (v/v) BSA (Sigma-Aldrich). Primary antibodies were incubated overnight at 4°C, while secondary antibodies were incubated at room temperature for 2 hours. After each antibody incubation, slides were washed 3x in PBS for 5 minutes. Slides were mounted in Fluoromount-G® (Southern Biotech, Sanbio, Uden, Netherlands). Primary and matching secondary antibodies are listed in Supplemental Table S2. Images were captured using the Zeiss LSM 880 confocal microscope or Leica DMI6000B high-content microscope.

#### Whole organoid staining

Whole mount staining was performed according to Takasato et al (Takasato et al., 2016). Briefly, organoids were fixed using 2% (w/v) PFA at 4°C for 20 minutes. After PBS wash, organoids were blocked in blocking buffer containing 10% (v/v) donkey serum (GeneTex, Irvine, California) and 0.6% (v/v) Triton-X in PBS at room temperature for 2h. Primary antibodies (Table S2) were diluted 1:300 and incubated at 4°C overnight. Next, organoids were washed using PBS-Triton-X (0.3% (v/v), PBTX). Secondary antibodies (Table S2) were diluted 1:400 in PBTX and incubated at room temperature for 2h. Organoids were washed using PBS and mounted using Fluoromount-G®. Images were captured using the Zeiss LSM 880 confocal microscope.

### Organoid filtration slit super resolution imaging

Sample processing and subsequent imaging was performed as described before (Artelt et al., 2018). In short, 4 μm paraffin sections were directly mounted on coverslips (VWR). To correct for PFA-induced autofluorescence, samples were incubated with 100 mM glycine in PBS for 10 minutes. Samples were blocked with 1% (v/v) fetal bovine serum, 1% (v/v) goat serum, 1% (v/v) bovine albumin and 0.1% (v/v) cold fish gelatin in PBS at RT for 1 hour. Primary antibody against nephrin (guinea pig, Progen GmbH, 1:100) was diluted in blocking solution and detected by a secondary anti-guinea pig Cy3-antibody (Jackson ImmunoResearch Laboratories Inc., 1:800) that was also diluted in blocking solution. 3D-SIM images were acquired using a Zeiss Elyra PS.1 system. Using Zeiss ZEN black software, 3D-SIM images were reconstructed. As control *in vivo* tissue we used a sections of Munich Wistar Frömter rats. The animals were housed under standard conditions. Animal procedures were approved by the German government officials (LANUV NRW 50.203.2 – AC 10/06) and performed in accordance with the European Communities Council Directive (86/609/EEC).

#### 2D podocyte staining

iPSC-derived podocytes and ciPODs cultured in 96-wells plates were fixed using using 2% (w/v) PFA for 10 minutes at RT. Cells were blocked using blocking buffer (same as used for whole organoid stain) at room temperature for 2h. Primary antibodies (Table S2) were diluted 1:100 and incubated at 4°C overnight. Next, cells were washed using PBTX and were incubated using secondary antibody diluted 1:200 in PBTX and incubated at room temperature for 2h. Cells were washed and mounted using Fluoromount-G®. Images were captured using the Leica DMI6000B high-content microscope.

### Protein harvesting, gel electrophoresis and Western blot

Tissue was collected and incubated on ice for 60 minutes using RIPA buffer containing 1X RIPA including a phosphatase inhibitor Na_3_VO_4_ (Cell Signaling, Leiden, the Netherlands), 1 mM phenylmethylsulfonyl fluoride (PMFS, Sigma) and 2% (w/v) sodium dodecyl sulphate (SDS, Sigma) in PBS. After incubation, tissue was centrifuged at 16,000 rcf for 15 minutes at 4°C. Supernatant containing protein was collected and stored at −80°C for further processing. Protein samples were mixed 1:4 with 4X Laemmli buffer (Bio-Rad, Lunteren, Netherlands) containing 10% (v/v) β-mercaptoethanol (Merck) and incubated at 95°C for 5 minutes. Samples were spun down and loaded onto a 4-20% (w/v) polyacrylamide precast mini-PROTEAN TGX gel (Bio-Rad) for subsequent gel-electrophoresis. Gels were then transferred onto a nitrocellulose blotting membrane (Amersham, GE Healtcare, Eindhoven, the Netherlands) and blocked for 1 hour at room temperature using Odyssey blocking buffer (LI-COR biosciences, Bio-Rad) in a 1:1 dilution with tris-buffered saline (TBS) containing 0.1% (v/v) Tween® 20 (Merck). Subsequently, membranes were washed 3 times for 5 minutes with TBS-T and incubated overnight with primary antibodies at 4°C. After washing the membranes again 3 times for 5 minutes with TBS-T, membranes were incubated with matching secondary antibodies for 1 h at room temperature. Membranes were washed using TBS and afterwards imaged and semi-quantified using a LI-COR Odyssey CLx imaging system. Primary- and matching secondary antibodies are listed in Supplemental Table S3.

### Transmission electron microscopy (TEM)

Organoids were fixed overnight using 2.5% (w/v) glutaraldehyde dissolved in 0.1 M sodium cacodylate. Tissue fragments were then fixed for 1 hour using 2% OsO_4_ buffered in pallade. Tissue was dehydrated through an increasing series of alcohol and embedded in Epon 812 (Sigma-Aldrich). Using a Leica Ultracut, 90 nm ultra-thin sections were cut, mounted on copper grids and contrasted at RT with uranyl acetate and lead citrate. Sections were assessed using a Jeol JEM 1400 electron microscope at 60kV. For image collection, a digital camera was used (Gatan Inc. Pleasanton, CA, USA).

### Fluorescence activated cell sorting (FACS)

Organoids cultured on Transwell™ plates were washed twice in PBS, dissociated using Accutase® and transferred to sterile 6-wells plates (without Transwell™ inserts). Organoids from similar culture conditions were pooled and incubated for 15 minutes at 37°C. During incubation, the cell suspension was resuspended every 2 minutes for homogenization purposes. After incubation, the cell suspension was put on ice for 1 minute to inactivate the Accutase®, cells were strained using a 40 μm cell strainer (Corning) and the cell strainer was washed repeatedly with 10% (v/v) FCS in DMEM/F12 to collect all cells. The strained cell suspension was centrifuged for at 250 rcf for 5 minutes at 4 °C and supernatant was discarded. Cells were washed with FACS buffer (PBS- 1% (v/v) BSA) once and incubated with primary antibodies for NPHS1 (1:100, RD systems, #AF4269) in FACS buffer for 45 minutes at 4 °C. After incubation, cells were centrifuged at 250 rcf for 5 minutes at 4°C and again washed with FACS buffer. After washing, cells were again centrifuged at 250 rcf for 5 minutes at 4°C and subsequently incubated with matching secondary antibodies (1:200, donkey anti-sheep Alexa Fluor™ 647, ThermoFisher, #A-21448) for 30 minutes at 4°C. Cells were centrifuged at 250 rcf for 5 minutes at 4°C and washed twice in FACS buffer. After washing, cells were incubated with DAPI (300 nM, ThermoFisher) in FACS buffer 10 minutes prior to FACS (for live/dead gating). Detection events were gated to select cells and doublet control was performed for both forward and side scatter. Living cells (DAPI^-^) were selected and gated on NPHS1^+^ (podocytes). Cells were sorted by immune fluorescence using a fluorescence cell sorter (BD Biosciences; FACSAria™ II SORP 18-color) and DIVA8 software. After cell sorting, DAPI^-^/NPHS1^+^ cells were collected in RLT buffer (RNeasy, Qiagen, Germantown, Maryland) for further processing.

### Plasma exposure and granule formation analysis using flow cytometry

D7+17 organoids were exposed to 10% (v/v) plasmapheresis plasma supplemented with anticoagulants heparin (100 μg/ml, Sigma Aldrich) and PPACK (10 μM, Santa Cruz Biotechnology) and incubated at 37°C, 5% (v/v) CO_2_, for 4 hours and 24 hours. The plasma was collected during plasmapheresis of a patient with posttransplant recurrent NS, during the active course of the disease and after remission. After collection, plasmapheresis plasma was aliquoted and stored at −80°C. As controls we used E6 culture medium, E6 medium supplemented with anti-coagulants heparin (100 μg/ml) and PPACK (10 μM) and plasmapheresis plasma collected from an iNS patient after kidney transplantation, with no disease recurrence (plasma presumably free of CPFs) and supplemented with anticoagulants. Next, single organoids were washed with PBS and digested using Accutase® for 15 minutes at 37°C, during this step the suspension was resuspended every 5 minutes. Accutase® was inactivated by adding DMEM/F12 supplemented with 10% (v/v) FBS. The organoid suspensions were centrifuged at 250 rcf for 5 minutes. Cells were washed once using wash buffer (PBS containing 1% (v/v) BSA) and again centrifuged. Next, the cells were incubated using a primary antibody against NPHS1 (R&D Systems, #AF4269), diluted 1:100 in FACS buffer (PBS supplemented with 1% (v/v) BSA) for 1 hour at 4°C. After 2 washing steps, the cells were incubated using a secondary antibody donkey anti-sheep Alexa Fluor 647™ (ThermoFisher, #A21448) 1:200 in FACS buffer for 45 min at 4°C. Cells were centrifuged at 250 rcf for 5 minutes at 4°C and washed twice in FACS buffer. After washing, cells were incubated with DAPI (300nM, ThermoFisher) in FACS buffer 10 minutes prior to FACS (for live/dead gating). Detection events were gated to select cells and doublet control was performed for both forward and side scatter. Living cells (DAPI^-^) were selected and gated on NPHS1^+^(podocytes). Cells were measured with a flow cytometer (NovoCyte 3000 flow cytometer, Agilent), as previously described by den Braanker *et al*. (den Braanker et al., 2021). The granule formation in living NPHS1+ podocytes was quantified by sideward scatter.

### Transcriptome analysis

#### Single-cell RNA sequencing

D7+17 organoids were washed with PBS and digested using Accutase® at 37°C for 15 minutes. Accutase® was inactivated by adding DMEM/F12 with 10% (v/v) FBS. The cell suspension was filtered through a 40 μm cell strainer and spun down at 1250 rcf for 5 minutes. To reduce background and obtain a single cell suspension, this step was repeated once more. Cells were then resuspended in PBS containing 0.04% (v/v) BSA and counted using a Neubauer counting chamber (Carl Roth, Karlsruhe, Germany). Next, 1000 cells/ μL per sample were loaded onto a 10x Chromium Next GEM Chip G according to the manufacturer’s instructions and processed in a Chromium controller (10X Genomics, Pleasanton, California). All single cell sequencing libraries were generated using 10x Chromium Next GEM v3.1 kits (10x Genomics) and sequencing was performed on a Novaseq6000 system (Illumina, San Diego, California), using an S2 v1.5 100 cycles flow cell (Illumina) with run settings 28-10-10-90 cycles. After sequencing, Cell Ranger (v.3.1.0) was used to align the reads to the human genome GRCh38, using default settings.

#### Standard single-cell RNA sequencing analysis work flow

The pre-processed reads were analyzed using the R package “Seurat” (version 4.0.3) (Hao et al., 2021). Data was loaded using the *Read10X* function and transformed into a Seurat object. A Seurat object was created with features (genes) that appeared in at least 3 cells and cells that contained at least 200 different features as a first filtering step. Next, cells were filtered out containing less than 200 or more than 3000 features to filter out damaged cells and doublets. Also, cells in which mitochondrial genes contributed for more than 20% to the total gene expression were believed to be damaged or apoptotic and therefore discarded. After these quality control checks, data was log normalized and variable features were identified using the *FindVariableFeatures* function with the vst selection method. Afterwards, the Seurat package was used to scale the data, and run a principal component analysis (PCA) dimensionality reduction. Data from the PCA was used to construct a Shared Nearest Neighbor (SNN) graph taking into account 20 principal components (PCs), which was subsequently used to identify clusters using the *FindClusters* function with the resolution set to 0.7. For visualization, the *RunUMAP* function was used. Next, clusters markers were obtained by running the *FindAllMarkers* function. Annotation of the clusters was performed manually using these cluster markers, and cluster markers described in literature and the Humphreys Kidney Interactive Transcriptomics (KIT) website (http://humphreyslab.com/SingleCell/).

#### Single-cell RNA sequencing in-depth podocyte cluster analysis

For in-depth podocyte analysis, the podocyte cluster was extracted from the total dataset and variable features were calculated again for this subset only. Next, the same workflow as described for the complete dataset after identifying variable features was performed. For comparative analysis with adult and fetal podocytes, datasets were used with gene expression omnibus accession number GSE118184 and GSE112570, respectively. Seurat objects were created from these datasets, dotplots for NPHS1, NPHS2, podocalyxin, synaptopodin and Wilms’ Tumor 1 were used to define the podocyte population within each dataset. This cluster was subsetted and merged with the subsetted podocytes from the organoids. Podocyte datasets were integrated using the *runHarmony* function from the Harmony (version 0.1.0) package (Korsunsky et al., 2019). Subsequently, we compared the podocyte clusters for marker expression using Seurat, pathway activity using PROGENy (version 1.14.0) and transcription factor activity using DoRothEA (version 1.4.1) and visualized using the pheatmap (version 1.0.12) R-package (Garcia-Alonso et al., 2019; Holland et al., 2020a; Holland et al., 2020b; Schubert et al., 2018). Cluster markers of podocyte clusters were determined using the *FindAllMarkers* function on a dataset with only these podocyte clusters (organoid, adult and fetal). These cluster markers were filtered to retain only significant markers (adjusted p value < 0.05) and were divided in upregulated (log fold change > 0) or downregulated (log fold change < 0). A list of significantly upregulated cluster markers was used to perform GO and KEGG enrichment analysis, using the clusterProfiler package (version 4.0.2) (Yu et al., 2012). MSigDB enrichment analysis was done using the ‘H’ or ‘C2’ category gene set from the msigdbr package (version 7.2.1) (Liberzon et al., 2015).

#### Ligand-receptor interaction analysis

In order to assess cellular interactions between cellular cluster within the organoid we replaced the podocyte cluster that was obtained by unsupervised clustering with the three podocyte subclusters that were identified during in-depth podocyte cluster analysis. Furthermore, cells positive for GATA3 and negative for CDH1 were named mesangial cells and cells positive for PECAM1 or CD34 or ICAM1 or CDH5 or PLVAP were named endothelial cells. Ligand-receptor analysis was performed with the addition of podocyte cluster 1, 2 and 3, mesangial cells and endothelial cells added to the original clusters identified by unsupervised clustering using Seurat. scRNA sequencing matrices were log-normalized and scaled and used as input for ligand receptor inference by CellphoneDB (CellphoneDB, version 2.0.5) (Efremova et al., 2020). Analysis was performed using the statistical_analysis mode of the CellphoneDB package. Using statistical significant interactions (p-value < 0.05) in the CellphoneDB output, ranking and visualization of the data was performed by CrossTalkeR (v 1.0.0) (Nagai et al., 2021). The same clusters were used to obtain GO, KEGG and Hallmark H enrichment in a similar way as described for the podocyte clusters (organoid, adult and fetal) described above.

#### Bulk RNA sequencing

Sequencing was performed at Single Cell Discoveries, a (single cell) sequencing provider located in the Netherlands using an adapted version of the CEL-seq protocol. In short, total RNA was extracted from FACS-sorted podocyte cell pellets using the Ambion PureLink RNA mini kit (ThermoFisher) according to the manufacturers protocol and used for library preparation and sequencing. mRNA was processed as described previously, following an adapted version of the single-cell mRNA seq protocol of CEL-Seq (Hashimshony et al., 2012; Simmini et al., 2014). Briefly, samples were barcoded with CEL-seq primers during a reverse transcription and pooled after second strand synthesis. The resulting cDNA was amplified with an overnight *in vitro* transcription reaction. From this amplified RNA, sequencing libraries were prepared with Illumina Truseq small RNA primers. Paired-end sequencing was performed on the Illumina Nextseq500 platform. Read 1 was used to identify the Illumina library index and CEL-Seq sample barcode. Read 2 was aligned to the hg19 human RefSeq transcriptome using BWA (Li and Durbin, 2010). Reads that mapped equally well to multiple locations were discarded. Mapping and generation of count Tables was done using the MapAndGo script1. Samples were normalized using RPM normalization.

#### Bulk RNA sequencing bioinformatic workflow

Bulk RNA sequencing data was used to produce heatmaps of known podocyte markers, pathway analysis using progeny and differential gene expression. Differential gene expression analysis was carried out using DESeq2 version 1.32.0 with internal statistical and normalization methods (i.e. multiple testing correction with Benjamini-Hochberg). The list of up- and downregulated genes that was obtained from DESeq2 for repaired podocytes versus mutant podocytes were used as input for enrichment analysis. Enrichment analysis for GO terms, KEGG pathways and Hallmark H gene sets was performed using clusterProfiler.

### RNAscope

Formalin-Fixed Paraffin-Embedded (FFPE) kidney organoid samples were pretreated according to the protocol of ACDbio (#322452, Bio-Techne). The probes that were used in this protocol were RNASCOPE® probe - Hs-COL4A3 (#461861, Bio-Techne), RNASCOPE® probe - Hs-PECAM1-O1-C2 (#487381-C2, Bio-Techne), RNASCOPE® probe - Hs-NPHS1-C3 (#416071-C3, Bio-Techne), and RNASCOPE® 3-Plex negative control probe (#320871, Bio-Techne). The RNAscope was performed according to the manufacturer’s protocol of ACDbio for fluorescent multiplex RNA *in situ* hybridization (#320293-USM). Images were captured using a Zeiss LSM 880 confocal microscope.

### Crispr/Cas NPHS2 repair c.413G>A to c.413G

Crispr Cas repair was performed by the Radboudumc Stem Cell Technology Center. In short, 70-90% confluent *NPHS2* mutant iPSCs were disassociated into single cells using TrypLE Select Enzyme for 2 minutes at 37°C. Cells were nucleofected with an RNP complex containing Alt-R® S.p. Cas9 Nuclease V3 (#1081058, IDT, Leuven, Belgium) coupled to Alt-R® S.p. CRISPR-Cas9 sgRNA 5’ AGAGTAATTATATTCCAACT 3’ (IDT) and Alt-R™ HDR donor oligo 5’ GTGTTTAGAAAAAAAAGAGTGTTTTTTTACCAGGGCCTTTGGCTCTTCCAGGAAGCAGATGTCCGAGTCGGAATATAATAACCCTTTCATACTCTTG 3’ (PS modification, antisense; IDT) using a 4D-nucleofector System X Unit (Lonza, Basel, Switzerland) and the P3 primary Cell 4D-Nucleofector® X kit (#V4XP-3024, Lonza). Nucleofected cells were seeded in Matrigel matrix-coated (1:50, Corning) 10 cm dishes and cultured in E8 medium supplemented with Alt-R® HDR Enhancer (#1081072, IDT) and E8 supplement for 24h. On day 2, medium was refreshed to E8 medium with E8 supplement and cells were cultured until colonies were observed. Multiple clones (~25) were picked, frozen and made into a pellet for subsequent processing. After proteinase K treatment, DNA was extracted from these cell pellets and, following PCR and purification, Sanger sequencing was performed to confirm successful gene editing. Clones were screened for correct gene editing using Chromas Lite and ICE analysis software (Synthego, Menlo park, CA). For sequencing data including the repaired mutation, see Supplemental Table S4.

### Fiji (ImageJ) quantification

Fluorescence images obtained from protamine sulfate experiments were analyzed using custom made macros in the free open-software Fiji/ImageJ (see Zenodo). First, the region of interest (ROI), was automatically determined through selection of NPHS1^+^ area (podocytes) using a readily available Fiji threshold algorithm. The same area selection algorithm was used for all raw images. Second, mean phalloidin intensity was measured in NPHS1^+^ areas to automatically determine changes in cytoskeleton arrangement (cytoplasmic contraction). As an internal quality control for automatic threshold selection, images containing the respective ROI selection were automatically created for post-processing analysis to ensure correct ROI selection in all conditions.

### Data and code availability

Single cell RNA sequencing data generated for this study is available from Gene Expression Omnibus (GEO (Barrett et al., 2013)) with GEO Series accession number GSE181954: https://www.ncbi.nlm.nih.gov/geo/query/acc.cgi?acc=GSE181954 and can be accessed with the secure token: gxkduoimbbafbab.

The scripts used for pathway analyses (PROGENy and DoRothEA), ligand-receptor analysis, as well as the Fiji macro for phalloidin cytoskeleton quantification are available on Zenodo: http://doi.org/10.5281/zenodo.5156000

### Statistical analysis

All data are expressed as mean ± SD of three independent experiments, unless stated otherwise. Statistical analysis was performed using one-way or two-way ANOVA analysis followed by Tukey post-test or, when appropriate, an unpaired *t* test with GraphPad Prism version 5.03 (La Jolla, CA). One asterisk was used to indicate significance with p<0.05, whereas two asterisks were used to indicate significance with p<0.01.

## References

Abrahamson, D.R., (2012). Role of the podocyte (and glomerular endothelium) in building the GBM. Semin Nephrol 32, 342–349.

Abrahamson, D.R., Hudson, B.G., Stroganova, L., Borza, D.B., St John, P.L., (2009). Cellular origins of type IV collagen networks in developing glomeruli. J Am Soc Nephrol 20, 1471–1479.

Adam, R.C., Yang, H., Ge, Y., Infarinato, N.R., Gur-Cohen, S., Miao, Y., Wang, P., Zhao, Y., Lu, C.P., Kim, J.E., Ko, J.Y., Paik, S.S., Gronostajski, R.M., Kim, J., Krueger, J.G., Zheng, D., Fuchs, E., (2020). NFI transcription factors provide chromatin access to maintain stem cell identity while preventing unintended lineage fate choices. Nat Cell Biol 22, 640–650.

Artelt, N., Siegerist, F., Ritter, A.M., Grisk, O., Schluter, R., Endlich, K., Endlich, N., (2018). Comparative Analysis of Podocyte Foot Process Morphology in Three Species by 3D Super-Resolution Microscopy. Front Med (Lausanne) 5, 292.

Barrett, T., Wilhite, S.E., Ledoux, P., Evangelista, C., Kim, I.F., Tomashevsky, M., Marshall, K.A., Phillippy, K.H., Sherman, P.M., Holko, M., Yefanov, A., Lee, H., Zhang, N., Robertson, C.L., Serova, N., Davis, S., Soboleva, A., (2013). NCBI GEO: archive for functional genomics data sets--update. Nucleic Acids Res 41, D991–995.

Chen, S., Kasama, Y., Lee, J.S., Jim, B., Marin, M., Ziyadeh, F.N., (2004). Podocyte-derived vascular endothelial growth factor mediates the stimulation of alpha3(IV) collagen production by transforming growth factor-beta1 in mouse podocytes. Diabetes 53, 2939–2949.

Colucci, M., Corpetti, G., Emma, F., Vivarelli, M., (2018). Immunology of idiopathic nephrotic syndrome. Pediatr Nephrol 33, 573–584.

den Braanker, D.J.W., Maas, R.J., Deegens, J.K., Yanginlar, C., Wetzels, J.F.M., van der Vlag, J., Nijenhuis, T., (2021). Novel in vitro assays to detect circulating permeability factor(s) in idiopathic focal segmental glomerulosclerosis. Nephrol Dial Transplant 36, 247–256.

Dong, L., Pietsch, S., Tan, Z., Perner, B., Sierig, R., Kruspe, D., Groth, M., Witzgall, R., Grone, H.J., Platzer, M., Englert, C., (2015). Integration of Cistromic and Transcriptomic Analyses Identifies Nphs2, Mafb, and Magi2 as Wilms’ Tumor 1 Target Genes in Podocyte Differentiation and Maintenance. J Am Soc Nephrol 26, 2118–2128.

Efremova, M., Vento-Tormo, M., Teichmann, S.A., Vento-Tormo, R., (2020). CellPhoneDB: inferring cell-cell communication from combined expression of multi-subunit ligand-receptor complexes. Nat Protoc 15, 1484–1506.

Eremina, V., Sood, M., Haigh, J., Nagy, A., Lajoie, G., Ferrara, N., Gerber, H.P., Kikkawa, Y., Miner, J.H., Quaggin, S.E., (2003). Glomerular-specific alterations of VEGF-A expression lead to distinct congenital and acquired renal diseases. J Clin Invest 111, 707–716.

Gallon, L., Leventhal, J., Skaro, A., Kanwar, Y., Alvarado, A., (2012). Resolution of recurrent focal segmental glomerulosclerosis after retransplantation. N Engl J Med 366, 1648–1649.

Garcia-Alonso, L., Holland, C.H., Ibrahim, M.M., Turei, D., Saez-Rodriguez, J., (2019). Benchmark and integration of resources for the estimation of human transcription factor activities. Genome Res 29, 1363–1375.

Garreta, E., Prado, P., Tarantino, C., Oria, R., Fanlo, L., Marti, E., Zalvidea, D., Trepat, X., Roca-Cusachs, P., Gavalda-Navarro, A., Cozzuto, L., Campistol, J.M., Izpisua Belmonte, J.C., Hurtado Del Pozo, C., Montserrat, N., (2019). Fine tuning the extracellular environment accelerates the derivation of kidney organoids from human pluripotent stem cells. Nat Mater 18, 397–405.

Geuens, T., Ruiter, F.A.A., Schumacher, A., Morgan, F.L.C., Rademakers, T., Wiersma, L.E., van den Berg, C.W., Rabelink, T.J., Baker, M.B., LaPointe, V.L.S., (2021). Thiol-ene cross-linked alginate hydrogel encapsulation modulates the extracellular matrix of kidney organoids by reducing abnormal type 1a1 collagen deposition. Biomaterials 275, 120976.

Guan, F., Villegas, G., Teichman, J., Mundel, P., Tufro, A., (2006). Autocrine VEGF-A system in podocytes regulates podocin and its interaction with CD2AP. Am J Physiol Renal Physiol 291, F422–428.

Hagmann, H., Brinkkoetter, P.T., (2018). Experimental Models to Study Podocyte Biology: Stock-Taking the Toolbox of Glomerular Research. Front Pediatr 6, 193.

Hale, L.J., Howden, S.E., Phipson, B., Lonsdale, A., Er, P.X., Ghobrial, I., Hosawi, S., Wilson, S., Lawlor, K.T., Khan, S., Oshlack, A., Quinlan, C., Lennon, R., Little, M.H., (2018). 3D organoid-derived human glomeruli for personalised podocyte disease modelling and drug screening. Nat Commun 9, 5167.

Hansen, K.U.I., Siegerist, F., Daniel, S., Schindler, M., Iervolino, A., Blumenthal, A., Daniel, C., Amann, K., Zhou, W., Endlich, K., Endlich, N., (2020). Prolonged podocyte depletion in larval zebrafish resembles mammalian focal and segmental glomerulosclerosis. FASEB J 34, 15961–15974.

Hao, Y., Hao, S., Andersen-Nissen, E., Mauck, W.M., 3rd, Zheng, S., Butler, A., Lee, M.J., Wilk, A. J., Darby, C., Zager, M., Hoffman, P., Stoeckius, M., Papalexi, E., Mimitou, E.P., Jain, J., Srivastava, A., Stuart, T., Fleming, L.M., Yeung, B., Rogers, A.J., McElrath, J.M., Blish, C.A., Gottardo, R., Smibert, P., Satija, R., (2021). Integrated analysis of multimodal single-cell data. Cell 184, 3573–3587 e3529.

Hashimshony, T., Wagner, F., Sher, N., Yanai, I., (2012). CEL-Seq: single-cell RNA-Seq by multiplexed linear amplification. Cell Rep 2, 666–673.

Hayashi, K., Sasamura, H., Nakamura, M., Azegami, T., Oguchi, H., Sakamaki, Y., Itoh, H., (2014). KLF4-dependent epigenetic remodeling modulates podocyte phenotypes and attenuates proteinuria. J Clin Invest 124, 2523–2537.

Holland, C.H., Szalai, B., Saez-Rodriguez, J., (2020a). Transfer of regulatory knowledge from human to mouse for functional genomics analysis. Biochim Biophys Acta Gene Regul Mech 1863, 194431.

Holland, C.H., Tanevski, J., Perales-Paton, J., Gleixner, J., Kumar, M.P., Mereu, E., Joughin, B. A., Stegle, O., Lauffenburger, D.A., Heyn, H., Szalai, B., Saez-Rodriguez, J., (2020b). Robustness and applicability of transcription factor and pathway analysis tools on single-cell RNA-seq data. Genome Biol 21, 36.

Homan, K.A., Gupta, N., Kroll, K.T., Kolesky, D.B., Skylar-Scott, M., Miyoshi, T., Mau, D., Valerius, M.T., Ferrante, T., Bonventre, J.V., Lewis, J.A., Morizane, R., (2019). Flow-enhanced vascularization and maturation of kidney organoids in vitro. Nat Methods 16, 255–262.

Jaffer, A.T., Ahmed, W.U., Raju, D.S., Jahan, P., (2011). Foothold of NPHS2 mutations in primary nephrotic syndrome. J Postgrad Med 57, 314–320.

Korsunsky, I., Millard, N., Fan, J., Slowikowski, K., Zhang, F., Wei, K., Baglaenko, Y., Brenner, M., Loh, P.R., Raychaudhuri, S., (2019). Fast, sensitive and accurate integration of single-cell data with Harmony. Nat Methods 16, 1289–1296.

Laboratory, T.H., Kidney Interactive Transcriptomics.

Li, H., Durbin, R., (2010). Fast and accurate long-read alignment with Burrows-Wheeler transform. Bioinformatics 26, 589–595.

Liberzon, A., Birger, C., Thorvaldsdottir, H., Ghandi, M., Mesirov, J.P., Tamayo, P., (2015). The Molecular Signatures Database (MSigDB) hallmark gene set collection. Cell Syst 1, 417–425.

Lu, Y., Ye, Y., Bao, W., Yang, Q., Wang, J., Liu, Z., Shi, S., (2017). Genome-wide identification of genes essential for podocyte cytoskeletons based on single-cell RNA sequencing. Kidney Int 92, 1119–1129.

Maas, R.J., Deegens, J.K., Smeets, B., Moeller, M.J., Wetzels, J.F., (2016). Minimal change disease and idiopathic FSGS: manifestations of the same disease. Nat Rev Nephrol 12, 768–776.

Maas, R.J., Deegens, J.K., Wetzels, J.F., (2014). Permeability factors in idiopathic nephrotic syndrome: historical perspectives and lessons for the future. Nephrol Dial Transplant 29, 2207–2216.

Munro, D.A.D., Wineberg, Y., Tarnick, J., Vink, C.S., Li, Z., Pridans, C., Dzierzak, E., Kalisky, T., Hohenstein, P., Davies, J.A., (2019). Macrophages restrict the nephrogenic field and promote endothelial connections during kidney development. Elife 8.

Musah, S., Mammoto, A., Ferrante, T.C., Jeanty, S.S.F., Hirano-Kobayashi, M., Mammoto, T., Roberts, K., Chung, S., Novak, R., Ingram, M., Fatanat-Didar, T., Koshy, S., Weaver, J.C., Church, G.M., Ingber, D.E., (2017). Mature induced-pluripotent-stem-cell-derived human podocytes reconstitute kidney glomerular-capillary-wall function on a chip. Nat Biomed Eng 1.

Nagai, J.S., Leimkuhler, N.B., Schaub, M.T., Schneider, R.K., Costa, I.G., (2021). CrossTalkeR: Analysis and Visualisation of Ligand Receptor Networks. Bioinformatics.

Nishibori, Y., Liu, L., Hosoyamada, M., Endou, H., Kudo, A., Takenaka, H., Higashihara, E., Bessho, F., Takahashi, S., Kershaw, D., Ruotsalainen, V., Tryggvason, K., Khoshnoodi, J., Yan, K., (2004). Disease-causing missense mutations in NPHS2 gene alter normal nephrin trafficking to the plasma membrane. Kidney Int 66, 1755–1765.

Noone, D.G., Iijima, K., Parekh, R., (2018). Idiopathic nephrotic syndrome in children. Lancet 392, 61–74.

Pages, G., Pouyssegur, J., (2005). Transcriptional regulation of the Vascular Endothelial Growth Factor gene--a concert of activating factors. Cardiovasc Res 65, 564–573.

Rauch, C., Feifel, E., Kern, G., Murphy, C., Meier, F., Parson, W., Beilmann, M., Jennings, P., Gstraunthaler, G., Wilmes, A., (2018). Differentiation of human iPSCs into functional podocytes. PLoS One 13, e0203869.

Reiser, J., Sever, S., Faul, C., (2014). Signal transduction in podocytes--spotlight on receptor tyrosine kinases. Nat Rev Nephrol 10, 104–115.

Rinschen, M.M., Godel, M., Grahammer, F., Zschiedrich, S., Helmstadter, M., Kretz, O., Zarei, M., Braun, D.A., Dittrich, S., Pahmeyer, C., Schroder, P., Teetzen, C., Gee, H., Daouk, G., Pohl, M., Kuhn, E., Schermer, B., Kuttner, V., Boerries, M., Busch, H., Schiffer, M., Bergmann, C., Kruger, M., Hildebrandt, F., Dengjel, J., Benzing, T., Huber, T.B., (2018). A Multi-layered Quantitative In Vivo Expression Atlas of the Podocyte Unravels Kidney Disease Candidate Genes. Cell Rep 23, 2495–2508.

Rinschen, M.M., Grahammer, F., Hoppe, A.K., Kohli, P., Hagmann, H., Kretz, O., Bertsch, S., Hohne, M., Gobel, H., Bartram, M.P., Gandhirajan, R.K., Kruger, M., Brinkkoetter, P.T., Huber, T.B., Kann, M., Wickstrom, S.A., Benzing, T., Schermer, B., (2017). YAP-mediated mechanotransduction determines the podocyte’s response to damage. Sci Signal 10.

Rudnicki, M., (2016). FSGS Recurrence in Adults after Renal Transplantation. Biomed Res Int 2016, 3295618.

Ryan, A.R., England, A.R., Chaney, C.P., Cowdin, M.A., Hiltabidle, M., Daniel, E., Gupta, A.K., Oxburgh, L., Carroll, T.J., Cleaver, O., (2021). Vascular deficiencies in renal organoids and ex vivo kidney organogenesis. Dev Biol 477, 98–116.

Sabo, A., Kress, T.R., Pelizzola, M., de Pretis, S., Gorski, M.M., Tesi, A., Morelli, M.J., Bora, P., Doni, M., Verrecchia, A., Tonelli, C., Faga, G., Bianchi, V., Ronchi, A., Low, D., Muller, H., Guccione, E., Campaner, S., Amati, B., (2014). Selective transcriptional regulation by Myc in cellular growth control and lymphomagenesis. Nature 511, 488–492.

Saleem, M.A., O’Hare, M.J., Reiser, J., Coward, R.J., Inward, C.D., Farren, T., Xing, C.Y., Ni, L., Mathieson, P.W., Mundel, P., (2002). A conditionally immortalized human podocyte cell line demonstrating nephrin and podocin expression. J Am Soc Nephrol 13, 630–638.

Schell, C., Wanner, N., Huber, T.B., (2014). Glomerular development--shaping the multi-cellular filtration unit. Semin Cell Dev Biol 36, 39–49.

Schubert, M., Klinger, B., Klunemann, M., Sieber, A., Uhlitz, F., Sauer, S., Garnett, M.J., Bluthgen, N., Saez-Rodriguez, J., (2018). Perturbation-response genes reveal signaling footprints in cancer gene expression. Nat Commun 9, 20.

Seckin, I., Uzunalan, M., Pekpak, M., Kokturk, S., Sonmez, H.A., Gungor, Z.B., Demirkiran, O.D., Saygi, H.I., Yaprak Sarac, E., (2017). The relationship between glomerular function and podocyte structure of pre-proteinuria and acute nephrosis in puromycin aminonucleoside-induced rat models: a comparative electron microscopic study. Rom J Morphol Embryol 58, 823–830.

Shabaka, A., Tato Ribera, A., Fernandez-Juarez, G., (2020). Focal Segmental Glomerulosclerosis: State-of-the-Art and Clinical Perspective. Nephron 144, 413–427.

Sharmin, S., Taguchi, A., Kaku, Y., Yoshimura, Y., Ohmori, T., Sakuma, T., Mukoyama, M., Yamamoto, T., Kurihara, H., Nishinakamura, R., (2016). Human Induced Pluripotent Stem Cell-Derived Podocytes Mature into Vascularized Glomeruli upon Experimental Transplantation. J Am Soc Nephrol 27, 1778–1791.

Shoji, J., Mii, A., Terasaki, M., Shimizu, A., (2020). Update on Recurrent Focal Segmental Glomerulosclerosis in Kidney Transplantation. Nephron 144 Suppl 1, 65–70.

Simmini, S., Bialecka, M., Huch, M., Kester, L., van de Wetering, M., Sato, T., Beck, F., van Oudenaarden, A., Clevers, H., Deschamps, J., (2014). Transformation of intestinal stem cells into gastric stem cells on loss of transcription factor Cdx2. Nat Commun 5, 5728.

Sison, K., Eremina, V., Baelde, H., Min, W., Hirashima, M., Fantus, I.G., Quaggin, S.E., (2010). Glomerular structure and function require paracrine, not autocrine, VEGF-VEGFR-2 signaling. J Am Soc Nephrol 21, 1691–1701.

Smeets, B., Kuppe, C., Sicking, E.M., Fuss, A., Jirak, P., van Kuppevelt, T.H., Endlich, K., Wetzels, J.F., Grone, H.J., Floege, J., Moeller, M.J., (2011). Parietal epithelial cells participate in the formation of sclerotic lesions in focal segmental glomerulosclerosis. J Am Soc Nephrol 22, 1262–1274.

Straatmann, C., Kallash, M., Killackey, M., Iorember, F., Aviles, D., Bamgbola, O., Carson, T., Florman, S., Vehaskari, M.V., (2014). Success with plasmapheresis treatment for recurrent focal segmental glomerulosclerosis in pediatric renal transplant recipients. Pediatr Transplant 18, 29–34.

Taguchi, A., Nishinakamura, R., (2017). Higher-Order Kidney Organogenesis from Pluripotent Stem Cells. Cell Stem Cell 21, 730–746 e736.

Takasato, M., Er, P.X., Chiu, H.S., Little, M.H., (2016). Generation of kidney organoids from human pluripotent stem cells. Nat Protoc 11, 1681–1692.

Takasato, M., Er, P.X., Chiu, H.S., Maier, B., Baillie, G.J., Ferguson, C., Parton, R.G., Wolvetang, E.J., Roost, M.S., Chuva de Sousa Lopes, S.M., Little, M.H., (2015). Kidney organoids from human iPS cells contain multiple lineages and model human nephrogenesis. Nature 526, 564–568.

Tanigawa, S., Islam, M., Sharmin, S., Naganuma, H., Yoshimura, Y., Haque, F., Era, T., Nakazato, H., Nakanishi, K., Sakuma, T., Yamamoto, T., Kurihara, H., Taguchi, A., Nishinakamura, R., (2018). Organoids from Nephrotic Disease-Derived iPSCs Identify Impaired NEPHRIN Localization and Slit Diaphragm Formation in Kidney Podocytes. Stem Cell Reports 11, 727–740.

Tory, K., Menyhard, D.K., Woerner, S., Nevo, F., Gribouval, O., Kerti, A., Straner, P., Arrondel, C., Huynh Cong, E., Tulassay, T., Mollet, G., Perczel, A., Antignac, C., (2014). Mutation-dependent recessive inheritance of NPHS2-associated steroid-resistant nephrotic syndrome. Nat Genet 46, 299–304.

Uchimura, K., Wu, H., Yoshimura, Y., Humphreys, B.D., (2020). Human Pluripotent Stem Cell-Derived Kidney Organoids with Improved Collecting Duct Maturation and Injury Modeling. Cell Rep 33, 108514.

van de Logt, A.E., Fresquet, M., Wetzels, J.F., Brenchley, P., (2019). The anti-PLA2R antibody in membranous nephropathy: what we know and what remains a decade after its discovery. Kidney Int 96, 1292–1302.

van den Berg, C.W., Koudijs, A., Ritsma, L., Rabelink, T.J., (2020). In Vivo Assessment of Size-Selective Glomerular Sieving in Transplanted Human Induced Pluripotent Stem Cell-Derived Kidney Organoids. J Am Soc Nephrol 31, 921–929.

van den Berg, C.W., Ritsma, L., Avramut, M.C., Wiersma, L.E., van den Berg, B.M., Leuning, D. G., Lievers, E., Koning, M., Vanslambrouck, J.M., Koster, A.J., Howden, S.E., Takasato, M., Little, M.H., Rabelink, T.J., (2018). Renal Subcapsular Transplantation of PSC-Derived Kidney Organoids Induces Neo-vasculogenesis and Significant Glomerular and Tubular Maturation In Vivo. Stem Cell Reports 10, 751–765.

van den Broek, M., Smeets, B., Schreuder, M.F., Jansen, J., (2021). The podocyte as a direct target of glucocorticoids in nephrotic syndrome. Nephrol Dial Transplant.

Veissi, S., Smeets, B., van den Heuvel, L.P., Schreuder, M.F., Jansen, J., (2020). Nephrotic syndrome in a dish: recent developments in modeling in vitro. Pediatr Nephrol 35, 1363–1372.

Verma, R., Venkatareddy, M., Kalinowski, A., Patel, S.R., Salant, D.J., Garg, P., (2016). Shp2 Associates with and Enhances Nephrin Tyrosine Phosphorylation and Is Necessary for Foot Process Spreading in Mouse Models of Podocyte Injury. Mol Cell Biol 36, 596–614.

Veron, D., Villegas, G., Aggarwal, P.K., Bertuccio, C., Jimenez, J., Velazquez, H., Reidy, K., Abrahamson, D.R., Moeckel, G., Kashgarian, M., Tufro, A., (2012). Acute podocyte vascular endothelial growth factor (VEGF-A) knockdown disrupts alphaVbeta3 integrin signaling in the glomerulus. PLoS One 7, e40589.

Warlich, E., Kuehle, J., Cantz, T., Brugman, M.H., Maetzig, T., Galla, M., Filipczyk, A.A., Halle, S., Klump, H., Scholer, H.R., Baum, C., Schroeder, T., Schambach, A., (2011). Lentiviral vector design and imaging approaches to visualize the early stages of cellular reprogramming. Mol Ther 19, 782–789.

Wharram, B.L., Goyal, M., Wiggins, J.E., Sanden, S.K., Hussain, S., Filipiak, W.E., Saunders, T.L., Dysko, R.C., Kohno, K., Holzman, L.B., Wiggins, R.C., (2005). Podocyte depletion causes glomerulosclerosis: diphtheria toxin-induced podocyte depletion in rats expressing human diphtheria toxin receptor transgene. J Am Soc Nephrol 16, 2941–2952.

Wu, H., Uchimura, K., Donnelly, E.L., Kirita, Y., Morris, S.A., Humphreys, B.D., (2018). Comparative Analysis and Refinement of Human PSC-Derived Kidney Organoid Differentiation with Single-Cell Transcriptomics. Cell Stem Cell 23, 869–881 e868.

Yamamoto-Nonaka, K., Koike, M., Asanuma, K., Takagi, M., Oliva Trejo, J.A., Seki, T., Hidaka, T., Ichimura, K., Sakai, T., Tada, N., Ueno, T., Uchiyama, Y., Tomino, Y., (2016). Cathepsin D in Podocytes Is Important in the Pathogenesis of Proteinuria and CKD. J Am Soc Nephrol 27, 2685–2700.

Yu, G., Wang, L.G., Han, Y., He, Q.Y., (2012). clusterProfiler: an R package for comparing biological themes among gene clusters. OMICS 16, 284–287.

Zhang, S.Y., Marlier, A., Gribouval, O., Gilbert, T., Heidet, L., Antignac, C., Gubler, M.C., (2004). In vivo expression of podocyte slit diaphragm-associated proteins in nephrotic patients with NPHS2 mutation. Kidney Int 66, 945–954.

