## Supplemental materials for "Human pluripotent stem cell-derived kidney organoids for personalized congenital and idiopathic nephrotic syndrome modeling"

Jitske Jansen[1,2,4,16], Bartholomeus T van den Berge[1,3,14], Martijn van den Broek[1,2,14], Rutger J Maas[3, 14], Brigith Willemsen[1], Christoph Kuppe[4,5], Katharina C Reimer[4,5,6], Gianluca Di Giovanni[1,2], Fieke Mooren[1], Quincy Nlandu[1], Helmer Mudde[1], Roy Wetzels[1], Dirk den Braanker[3], Naomi Parr[3], James S Nagai[7,8], Vedran Drenic[9], Ivan G Costa[7,8], Eric Steenbergen[1], Tom Nijenhuis[3], Nicole Endlich[9,10], Nicole CAJ van de Kar[2], Rebekka K Schneider[6,11,12], Jack FM Wetzels[3], Johan van der Vlag[3], Rafael Kramann[4,5,13], Michiel F Schreuder[2,15], Bart Smeets[1, 15, 16]

[1]Department of Pathology, Radboud Institute for Molecular Life Sciences, Radboud university medical center, P.O. Box 9101, 6500 HB Nijmegen, the Netherlands
[2]Department of Pediatric Nephrology, Radboud Institute for Molecular Life Sciences, Radboud University Medical Center, Amalia Children's Hospital, P.O. Box 9101, 6500 HB Nijmegen, the Netherlands
[3]Department of Nephrology, Radboud Institute for Molecular Life Sciences, Radboud Institute for Health Sciences, Radboud University Medical Center, P.O. Box 9101, 6500 HB Nijmegen, The Netherlands
[4]Institute of Experimental Medicine and Systems Biology, Medical Faculty RWTH Aachen University, Pauwelsstrasse 30, 52074 Aachen, Germany
[5]Division of Nephrology and Clinical Immunology, RWTH Aachen University, Aachen, Germany
[6]Institute for Biomedical Technologies, Department of Cell Biology, RWTH Aachen University, Aachen, Germany
[7]Institute for Computational Genomics, University Hospital RWTH Aachen, Germany
[8]Joint Research Center for Computational Biomedicine, RWTH Aachen University Hospital, Aachen, Germany
[9]NIPOKA GmbH, 17489 Greifswald, Germany
[10]Department of Anatomy and Cell Biology, University Medicine Greifswald, 17489 Greifswald, Germany
[11]Department of Developmental Biology, Erasmus Medical Center, Rotterdam, The Netherlands
[12]Oncode Institute, Erasmus Medical Center, Rotterdam, The Netherlands
[13]Department of Internal Medicine, Nephrology and Transplantation, Erasmus Medical Center, Rotterdam, NL
[14]These authors contributed equally
[15]These authors contributed equally
[16]Lead contact

<u>Corresponding authors:</u>
Bart Smeets, PhD
Jitske Jansen, PhD
Radboud University Medical Center
Radboud Institute for Molecular Life Sciences

Department of Pathology
P.O. Box 9101
6500 HB Nijmegen, the Netherlands



**Supplemental Table S1.** Top 20 DE genes based on adjusted p-value

| | PT/LH/DT | Podocyte precursor | Podocyte subcluster 1 | Podocyte subcluster 2 | Podocyte subcluster 3 | Loop progenitor | DT/CD | EC precursor | Mesengial precursor | Stroma 1 | Stroma 2 | Stroma 3 | Stroma 4 | Stroma 5 | Stroma/neural progenitor | Stroma/ neuron | Neuron/ glial |
|---|---|---|---|---|---|---|---|---|---|---|---|---|---|---|---|---|---|
| 1 | IRX1 | WT1 | NPHS2 | CLIC5 | ITGA3 | KIF20A | LBX1 | CD34 | GATA3 | CTSC | CD24 | ADCYAP1 | ISL1 | MEOX2 | CD81 | CLSPN | ZIC2 |
| 2 | IRX2 | MAFB | SOST | PTPRO | PTPRO | DLGAP5 | PITX2 | PLVAP | NEFM | COL3A1 | ALX4 | CYTL1 | OSR2 | DLX2 | COL3A1 | XRCC2 | SOX2 |
| 3 | MAB21L2 | GADD45A | TCF21 | SOST | PODXL | CENPA | POU4F1 | EBF3 | ANGPT1 | SULT1E1 | PRRX1 | CNTN3 | NEFM | DLX1 | COL1A2 | TYMS | PAX3 |
| 4 | RFLNA | NPHS2 | PODXL | WT1 | NPHS1 | CENPE | TMEM108 | VCAM1 | FLRT3 | CRABP1 | COL21A1 | PRSS12 | SIM2 | RALYL | MT-ND1 | UHRF1 | RFX4 |
| 5 | CLDN11 | MDK | PTPRO | SBSPON | PLA2R1 | KNL1 | GATA3 | CRABP1 | VCAN | CHST2 | NOVA1 | TAC1 | ZNF385D | PALMD | HNF1B | CENPU | CACNG8 |
| 6 | MECOM | MT-CO2 | CLIC5 | PODXL | ENPEP | CDC20 | GRIA4 | IFITM3 | FN1 | NAV3 | CXCL14 | VEGFC | HAND2 | FSTL5 | LAPTM4A | DTL | POU3F2 |
| 7 | CXCL12 | MARCKS | WT1 | NPHS2 | ST3GAL6 | GTSE1 | MYF6 | COL12A1 | PDGFC | STMN2 | MAB21L2 | CRABP1 | SLC26A7 | MGP | SFRP2 | FAM111B | ZIC5 |
| 8 | PTN | FTH1 | DDN | NPHS1 | NPHS2 | HMMR | LYPD1 | COL6A3 | PTH1R | S100A10 | MAB21L1 | MEIS2 | CHRM2 | TNMD | HSP90B1 | ORC6 | CHL1 |
| 9 | SIX2 | MT-ATP6 | AQP3 | AQP3 | AC008264.2 | TACC3 | CAMK2N1 | LAMA4 | FHL1 | NDST3 | ALDH1A2 | TMSB4X | TTN | IFI44L | GAS1 | RAD51AP1 | SMOC1 |
| 10 | USH1C | KCNQ1OT1 | ENPEP | TCF21 | CLIC5 | PLK1 | GATA3-AS1 | CD99 | AC060834.2 | PENK | POSTN | FAT4 | GUCY1A1 | HPGD | FBLN1 | MYBL2 | MSX1 |
| 11 | COL2A1 | MT-CO3 | SBSPON | DDN | SPOCK2 | NUF2 | OLFML2B | PCOLCE | TNMD | MEIS2 | ALX1 | CDH2 | CPE | TRPS1 | PDGFRA | PCLAF | PHYHIPL |
| 12 | COL9A2 | SOX4 | PLA2R1 | VAMP8 | SBSPON | ASPM | PDGFC | HMGA2 | MEOX2 | BCHE | CSRP2 | MIR137HG | NRP2 | MKX | MT-CYB | GINS2 | PI15 |
| 13 | EGFL6 | MT-ND4 | ST3GAL6 | ST3GAL6 | SOST | CCNB2 | LIN7A | USH1C | TMSB4X | ACKR3 | LVRN | SPOCK3 | TBX2 | COL11A1 | CPE | ATAD5 | MAP6 |
| 14 | ERG | MT-ND1 | SPOCK2 | ENPEP | VEGFA | CKAP2L | SNCA | DDR2 | FAM110B | TPM1 | FAM196A | USH1C | SORCS1 | FIBIN | MT-CO3 | E2F1 | LMX1A |
| 15 | PLD5 | MT-CO1 | VAMP8 | MPP5 | WT1 | KIF4A | FLRT3 | BTBD3 | BCHE | TWIST1 | LUM | RPL3 | CCDC141 | CSMD3 | MDK | MCM10 | TFAP2B |
| 16 | TSPAN13 | CRABP1 | CLDN5 | CLDN5 | PRODH2 | TROAP | SHISA2 | SERPINH1 | MDK | COL6A3 | PRRX2 | RPL10 | NRXN1 | C3orf80 | CALR | CDC45 | GDF7 |
| 17 | MIA | ANXA1 | NPHS1 | SPOCK2 | MPP5 | TOP2A | PTPRZ1 | ROBO2 | NR2F1 | KLHL4 | SOX11 | RPL13 | IGFBP3 | RGCC | MT-ATP6 | CENPK | RAB3C |
| 18 | CYP1B1 | AIF1 | MRGPRF | ARHGAP29 | PTPRQ | MKI67 | EGFLAM | TPM1 | KIF26B | PARM1 | TBX3 | HGF | COL21A1 | THBS4 | IL11RA | CDC6 | SATB2 |
| 19 | SOX5 | TPM2 | STON2 | PLA2R1 | PLTP | KIF2C | CRABP1 | CHST2 | ADAMTS6 | SLIT2 | PCDH9 | RPL10A | SIM1 | RUNX1T1 | FSTL1 | MCM4 | TTYH1 |
| 20 | FAT3 | COL3A1 | TPPP3 | GADD45A | MAGI2 | BIRC5 | ADAMTS6 | EDNRA | CREB5 | MAB21L2 | PCDH10 | RPL30 | PDZRN3 | PTGFR | APP | FANCD2 | WSCD2 |

PT: proximal tubule, LH: loop of Henle, DT: distal tubule, CD: collecting duct, EC: endothelial cells.

**Supplemental Table S2.** Antibodies used for immunofluorescence staining.

| Primary antibody | Secondary antibody |
|---|---|
| Human Nephrin Antibody (AF4269, R&D Systems) | Donkey anti-sheep Alexa Fluor™ 647 (A21448, Thermo Fisher) |
| Anti-Podocin antibody (P0372, Sigma Aldrich) | Donkey anti-rabbit Alexa Fluor™ 568 (A10042, Thermo Fisher) |
| Anti-Human WT1 clone 6F-H2 (M3561, Dako) | Donkey anti-mouse Alexa Fluor™ 488 (A21202, Thermo Fisher) |
| Anti-synaptopodin antibody (65194, Progen) | Donkey anti-mouse Alexa Fluor™ 488 (A21202, Thermo Fisher) |
| Monoclonal Anti-PLA2R antibody (AMAB90772, Sigma Aldrich) | Donkey anti-mouse Alexa Fluor™ 488 (A21202, Thermo Fisher) |
| Human CD31/PECAM-1 Antibody (BBA7, R&D Systems) | Donkey anti-mouse Alexa Fluor™ 488 (A21202, Thermo Fisher) |
| PV-1 (PLVAP) antibody (HPA002279, Sigma-Aldrich) | Donkey anti-rabbit Alexa Fluor™ 568 (A10042, Thermo Fisher) |
| VE-cadherin (F-8) AC (sc-9989 AC, Santa Cruz Biotechnology) | Donkey anti-mouse Alexa Fluor™ 647 (A31571, Thermo Fisher) |
| Lotus Tetragonolobus Lectin (LTL) biotinylated (B-1325, Vector Laboratories) | Streptavidin, Alexa Fluor™ 405 conjugate (S32351, Thermo Fisher) |
| Anti-E-Cadherin clone 36 (610181, BD Biosciences) | Donkey anti-mouse Alexa Fluor™ 488 (A21202, Thermo Fisher) |
| GATA-3 (D13C9) monoclonal Antibody (5852, Cell Signaling) | Donkey anti-rabbit Alexa Fluor™ 647 (A31573, Thermo Fisher) |
| Flash Phalloidin™ Green 488 (424201, Biolegend) | n.a. |
| Anti-EHD3 antibody (HPA049890, Sigma-Aldrich) | Donkey anti-rabbit Alexa Fluor™ 568 (A10042, Thermo Fisher) |

**Supplemental Table S3.** Antibodies used for western blotting.

| **Primary antibody** | **Secondary antibody** |
|---|---|
| Anti-Nephrin (phosphor Y1176 + Y1193) Antibody (ab80299, Abcam) | Goat anti-rabbit, Alexa Fluor™ 680 (A-21076, Thermo Fisher) |
| Anti-Nephrin antibody (sc-19000, Santa Cruz) | Donkey anti-goat, Alexa Fluor™ 680 (A-21084, Thermo Fisher) |
| Anti-γ-Tubulin antibody (T6557, Sigma) | Donkey anti-mouse, DyLight 800 (SA5-10172, Thermo Fisher) |

**Supplemental Table S4.** iPS-NPHS2 mutant CRISPR sequencing data

| |
|---|
| **Sequencing data before CRISPR repair (NPHS2 exon 3):**<br>ATCTAGAAAAAAAAAAAATCCAATTGAATCATTTTGAAGCAGCCTCAGAAGAAATTGCACTCTGA<br>AACAAAACATCACAAGTGGTATTAAAATATACTCCCTCTTTCCATTAAATAGCTAATGAGCAATA<br>CATATAAAAGCTAGTGCAGAACTCACAGTAAAATATAAGATTTAATATGCCTTGATAAGATTAA<br>TTTAGGGAAAGTTGGCCATGGATTTTAGATAATCATAAGTCTTTAATCAAAATTCTGTCTATGGG<br>TTCAAAAATTAACATGGTTAATATACTTTTTCATTTCTGAAATTTTACACTTACTAAATATAGATT<br>TTGGAAACTTAAGTATTAATAGAAATTTTTTCCTGGTTCTCAAAACAAAAAATTTCTGATATCTA<br>GGATCATTCTTATGCCAAGGCCTTTTGAAGACTTTTTCTTTCTGGGAGTGATTTGAAAGGATTAA<br>ATTTCTCTTTAGGTTGTA*CAAGAGTATGAAAGAGTAATTATATTCC*__A__*ACTGGGACATCTGCTTCCTGGA*<br>*AGAGCCAAAGGCCCTGGTAAAAAAACACTCTTTTTTTTCTAAACAC*CTCTCTCCTGACTTGCCAATTTC<br>TTCAACCCATGCAGATTTGTAATATGGACCTCAGATTAAATGAAGTAACTTGATTCATGATATCT<br>GAATTTTCCAATCTGTTACTTATAGGTTATTCAAATATTCTTCAGAGACTATTACTACTAGGTCAT<br>AGGTAGCCAAGAGAGAGAATTGGTACAGAGAGCCCACATGCCAGGGCAAGGCTTGCTGGAATA<br>GCAAGTTAGCTTAGGACCAATGGCTGGGGACTGATTTGAGTACGATGTGTATATGGTCAGAGTC<br>CTGTGAATTCTCAGAGAAAAGCTGAGCTAGTTCCCACTCCAGGTGCTCAGATCCAATCATGAGA<br>AACAGGCAAAGTTCAGCATTCAAATAACAAGTTGCTCTTCAGTTATAGGGATTTAAAAAACATA<br>GATTAAGCATTTCTTGTGGCTCA |
| **Sequencing data after CRISPR repair (NPHS2 exon 3):**<br>ATCTAGAAAAAAAAAAAATCCAATTGAATCATTTTGAAGCAGCCTCAGAAGAAATTGCACTCTGA<br>AACAAAACATCACAAGTGGTATTAAAATATACTCCCTCTTTCCATTAAATAGCTAATGAGCAATA<br>CATATAAAAGCTAGTGCAGAACTCACAGTAAAATATAAGATTTAATATGCCTTGATAAGATTAA<br>TTTAGGGAAAGTTGGCCATGGATTTTAGATAATCATAAGTCTTTAATCAAAATTCTGTCTATGGG<br>TTCAAAAATTAACATGGTTAATATACTTTTTCATTTCTGAAATTTTACACTTACTAAATATAGATT<br>TTGGAAACTTAAGTATTAATAGAAATTTTTTCCTGGTTCTCAAAACAAAAAATTTCTGATATCTA<br>GGATCATTCTTATGCCAAGGCCTTTTGAAGACTTTTTCTTTCTGGGAGTGATTTGAAAGGATTAA<br>ATTTCTCTTTAGGTTGTA*CAAGAGTATGAAAGGGTTATTATATTCC*__*G*__*ACTGGGACATCTGCTTCCTGGA*<br>*AGAGCCAAAGGCCCTGGTAAAAAAACACTCTTTTTTTTCTAAACAC*CTCTCTCCTGACTTGCCAATTTC<br>TTCAACCCATGCAGATTTGTAATATGGACCTCAGATTAAATGAAGTAACTTGATTCATGATATCT<br>GAATTTTCCAATCTGTTACTTATAGGTTATTCAAATATTCTTCAGAGACTATTACTACTAGGTCAT<br>AGGTAGCCAAGAGAGAGAATTGGTACAGAGAGCCCACATGCCAGGGCAAGGCTTGCTGGAATA<br>GCAAGTTAGCTTAGGACCAATGGCTGGGGACTGATTTGAGTACGATGTGTATATGGTCAGAGTC<br>CTGTGAATTCTCAGAGAAAAGCTGAGCTAGTTCCCACTCCAGGTGCTCAGATCCAATCATGAGA<br>AACAGGCAAAGTTCAGCATTCAAATAACAAGTTGCTCTTCAGTTATAGGGATTTAAAAAACATA<br>GATTAAGCATTTCTTGTGGCTCA |

RED = sgRNA, *Cursive = Homology Directed Repair (HDR) template,* **BOLD = mutation (edit)**

The HDR Template contains silent mutations in the protospacer adjacent motif (PAM) region in order to prevent re-cutting after successful editing.

Original sequence:
CAAGAGTATGAAAGAGTAATTATATTCCAACTGGGACATCTGCTTCCTGGAAGAGCCAAAGGCCCT
GGTAAAAAAACACTCTTTTTTTTCTAAACAC

Sequence after successful editing and incorporation of silent mutations:
CAAGAGTATGAAAGGGTTATTATATTCCGACTGGGACATCTGCTTCCTGGAAGAGCCAAAGGCCCT
GGTAAAAAAACACTCTTTTTTTTCTAAACAC

**Supplemental Table S5.** Reagent and resource table

| Reagent or Resource | Source | Identifier |
|---|---|---|
| **Antibodies** | | |
| anti-human nephrin, polyclonal sheep IgG | R&D Systems | Cat#AF4269 |
| anti-human nephrin, polyclonal guinea pig | Progen | Cat#GP-N2 |
| anti-human podocin, polyclonal rabbit IgG | Sigma Aldrich | Cat#P0372 |
| anti-human WT1, monoclonal mouse IgG1 | Dako | Cat#M3561 |
| anti-human synaptopodin, monoclonal mouse IgG1 | Progen Biotechnik | Cat#65194 |
| anti-human PLA2R, monoclonal mouse IgG1 | Sigma Aldrich | Cat#AMAB90772 |
| anti-human CD31/PECAM-1, monoclonal mouse IgG1 | R&D Systems | Cat#BBA7 |
| anti-human PLVAP, polyclonal rabbit IgG | Sigma Aldrich | Cat#HPA002279 |
| anti-human VE-cadherin, monoclonal mouse IgG1 | Santa Cruz Biotechnology | Cat#sc-9989 |
| biotinylated lotus tetragonolobus lectin (LTL) | Vector Laboratories | Cat#B1325 |
| anti-human E-cadherin, monoclonal mouse IgG2a | BD Biosciences | Cat#610181 |
| anti-human GATA-3, monoclonal rabbit IgG | Cell Signaling | Cat#5852 |
| Flash Phalloidin™ Green 488 | Biolegend | Cat#424201 |
| anti-human EHD3, polyclonal rabbit IgG | Sigma Aldrich | Cat#HPA049890 |
| anti-human nephrin (phospho Y1176/Y1193), monoclonal rabbit IgG | Abcam | Cat#ab80299 |
| anti-human nephrin, polyclonal goat IgG | Santa Cruz Biotechnology | Cat#sc-1900 |
| anti-human γ-tubulin, monoclonal mouse IgG | Sigma Aldrich | Cat#T6557 |
| Alexa Fluor donkey anti-sheep IgG 647 (H+L) | Thermo Fisher | Cat#A21448 |
| AffinuPure anti-guinea pig IgG Cy3 (H+L) | Jackson ImmunoResearch Laboratories Inc. | Cat#706-165-148 |
| Alexa Fluor donkey anti-rabbit IgG 647 (H+L) | Thermo Fisher | Cat#A31573 |
| Alexa Fluor donkey anti-rabbit IgG 568 (H+L) | Thermo Fisher | Cat#A10042 |
| Alexa Fluor donkey anti-mouse IgG 647 (H+L) | Thermo Fisher | Cat#A31571 |
| Alexa Fluor donkey anti-mouse IgG 488 (H+L) | Thermo Fisher | Cat#A21202 |
| Streptavidin Alexa Fluor 405 conjugate | Thermo Fisher | Cat#S32351 |

| | | |
|---|---|---|
| Alexa Fluor goat anti-rabbit IgG 680 | Thermo Fisher | Cat#A-21076 |
| Alexa Fluor donkey anti-goat IgG 680 | Thermo Fisher | Cat#A-21084 |
| Donkey anti-mouse IgG DyLight 800 (H+L) | Thermo Fisher | Cat#SA5-10172 |
| **Chemicals, Peptides, and Recombinant Proteins** | | |
| Essential 8 FLEX Medium | Thermo Fisher | Cat#A2858501 |
| Essential 6 Medium | Thermo Fisher | Cat#A1516401 |
| UltraPure EDTA (0.5M) pH 8.0 | Thermo Fisher | Cat#15575020 |
| Geltrex LDEV-free, hESC-Qualified, reduced growth factor basement membrane matrix | Thermo Fisher | Cat#A1413302 |
| Antibiotic-Antimycotic | Thermo Fisher | Cat#15240062 |
| CHIR 99021 | R&D Systems | Cat#4423/10 |
| Recombinant Human FGF-9 Protein | R&D Systems | Cat#273-F9-025 |
| Heparin sodium salt from porcine intestinal mucosa | Sigma Aldrich | Cat#H4784 |
| Recombinant Human BMP-7 Protein | R&D Systems | Cat#354-BP-010 |
| Human Activin A | Milteyni Biotec | Cat#130-115-010 |
| Retinoic acid | Sigma Aldrich | Cat#2625-100MG |
| Human $VEGF_{165}$ | R&D Systems | Cat#293-VE-010/CF |
| MEM Non-Essential Amino Acids Solution | Thermo Fisher | Cat#11140050 |
| TrypLE Select Enzyme, no phenol red | Thermo Fisher | Cat#12563-029 |
| RevitaCell Supplement | Thermo Fisher | Cat#A2644501 |
| Trypsin-EDTA (0.05%), phenol red | Thermo Fisher | Cat#25300054 |
| Corning® Transwell® polyester membrane cell culture inserts, 24 mm Transwell with 0.4 µm pore polyester membrane insert, TC-treated, sterile | Sigma Aldrich | Cat#CLS3450 |
| Corning® Falcon® 96-well, clear flat bottom, TC-treated, sterile imaging microplate | Sigma Aldrich | Cat#353219 |
| DMEM/F-12 | Thermo Fisher | Cat#11320074 |
| Human EGF | Sigma Aldrich | Cat#E9644 |
| Protamine sulfate | Sigma Aldrich | Cat#P3369 |
| Insulin-transferrin-sodium selenite media supplement | Sigma Aldrich | Cat#I1884-1VL |
| Penicillin-Streptomycin (5000 U/mL) | Thermo Fisher | Cat#15070063 |

| | | |
|---|---|---|
| Fetal Bovine Serum, heat inactivated | Thermo Fisher | Cat#16140071 |
| Accutase® solution | Sigma Aldrich | Cat#A6964 |
| PPACK dihydrochloride | Santa Cruz Biotechnology | Cat#sc-201291 |
| **Critical Commercial Assays** | | |
| PureLink RNA mini kit | Thermo Fisher | Cat#12183018A |
| PSC Cryopreservation Kit | Thermo Fisher | Cat#351-FS-050 |
| RNAscope® Multiplex Fluorescent Reagent Kit v2 assay | Advanced Cell Diagnostics | Cat#323100 |
| RNAscope® probe - Hs-COL4A3 | Advanced Cell Diagnostics | Cat#461861 |
| RNAscope® probe - Hs-PECAM1-O1-C2 | Advanced Cell Diagnostics | Cat#487381-C2 |
| RNAscope® probe - Hs-NPHS1-C3 | Advanced Cell Diagnostics | Cat#416071-C3 |
| RNAscope® 3-Plex negative control probe | Advanced Cell Diagnostics | Cat#320871 |
| Chromium Next GEM Single Cell 3' Kit v3.1,<br><br>16 rxns | 10x Genomics | Cat#1000268 |
| Dual Index Kit TT Set A, 96 rxns | 10x Genomics | Cat#1000215 |
| Chromium Next GEM Chip G Single Cell Kit,<br><br>16 rxns | 10x Genomics | Cat#1000127 |
| Library Construction Kit 16 rxns | 10x Genomics | Cat#1000190 |
| NovaSeq 6000 S2 Reagent Kit v1.5 (100 cycles) | Illumina | Cat#20028316 |
| **Deposited Data** | | |
| scRNAseq data | This paper; deposited on Gene Expression Omnibus (GEO) | GSE181954: https://www.ncbi.nlm.nih.gov/geo/query/acc.cgi?acc=GSE181954 |
| Scripts and codes for data analysis | This paper; deposited on Zenodo | http://doi.org/10.5281/zenodo.5156000 |
| **Experimental Models: Cell Lines** | | |

| | | |
|---|---|---|
| Human induced pluripotent stem cell line iPS 15 | SCTC Radboud UMC, The Netherlands | iPS 15 |
| Human induced pluripotent stem cell line iPS 134 | iPS core facility Erasmus MC, The Netherlands | iPS 134 |
| Human induced pluripotent stem cell line iPS 19 | SCTC Radboud UMC, The Netherlands | iPS19_91 |
| Conditionally immortalized human podocyte cell line | Prof. Saleem, Bristol, UK | ciPOD |
| **Software and Algorithms** | | |
| ImageJ version Fiji 1.51n | National Institutes of Health, USA | https://imagej.nih.gov/ij/ |
| ZEN 3.0 blue edition | Carl Zeiss | https://www.zeiss.de/mikroskopie/produkte/mikroskopsoftware/zen.html |
| Cell Ranger version 3.1.0 | 10x genomics | https://support.10xgenomics.com/single-cell-gene-expression/software/release-notes/2-1 |
| Seurat version 4.0.3 | Hao and Hao et al. Integrated analysis of multimodal single-cell data. Cell (2021) | https://satijalab.org/seurat/ |
| R 4.1.0 | CRAN | https://www.R-project.org |
| Adobe InDesign CC2021 | Adobe Systems Inc. | https://www.adobe.com/nl/products/indesign.html |
| Adobe Illustrator CC 2021 | Adobe Systems Inc. | RRID:SCR_010279 |
| Adobe Photoshop CC 2021 | Adobe Systems Inc. | RRID:SCR_014199 |
| GraphPad Prism version 9.0 | GraphPad Software Inc. | RRID:SCR_002798 |

| | | |
|---|---|---|
| clusterProfiler 4.0.2 | Wu T, Hu E, Xu S, Chen M, Guo P, Dai Z, Feng T, Zhou L, Tang W, Zhan L, Fu x, Liu S, Bo X, Yu G (2021). "clusterProfiler 4.0: A universal enrichment tool for interpreting omics data." The Innovation, 2(3), 100141. doi: 10.1016/j.xinn.2021.100141. | https://bioconductor.org/packages/release/bioc/html/clusterProfiler.html |
| Harmony 0.1.0 | Korsunsky, I., Millard, N., Fan, J. et al. Fast, sensitive and accurate integration of single-cell data with Harmony. Nat Methods 16, 1289–1296 (2019). https://doi.org/10.1038/s41592-019-0619-0 | https://portals.broadinstitute.org/harmony/articles/quickstart.html#session-info-1 |
| PROGENy 1.14.0 | Schubert, M., Klinger, B., Klünemann, M. *et al.* Perturbation-response genes reveal signaling footprints in cancer gene expression. *Nat Commun* **9,** 20 (2018). https://doi.org/10.1038/s41467-017-02391-6 | https://saezlab.github.io/progeny/ |
| DoRothEA 1.4.1 | Garcia-Alonso L, Iorio F, Matchan A, et al. Transcription Factor Activities Enhance Markers of Drug Sensitivity in Cancer. Cancer Research. 2018 Feb;78(3):769-780. DOI: 10.1158/0008-5472.can-17-1679. | https://saezlab.github.io/dorothea/index.html |

| | | |
|---|---|---|
| Pheatmap 1.0.12 | Raivo Kolde (2019). Pheatmap: Pretty heatmaps. | https://CRAN.R-project.org/package=pheatmap |
| CrossTalkeR 1.0.2 | CrossTalkeR: Analysis and Visualisation of Ligand Receptor Networks<br><br>James S. Nagai, Nils B. Leimkühler, Rebekka K. Schneider, Ivan G.Costa<br><br>bioRxiv 2021.01.20.427390; doi: https://doi.org/10.1101/2021.01.20.427390 | https://github.com/<br><br>CostaLab/CrossTa<br><br>lkeR |
| CellPhoneDB 2.0.5 | Single-cell reconstruction of the early maternal-fetal interface in humans. Vento-Tormo R, Efremova M, et al., Nature. 2018 Nov;563(7731):347-353. doi: 10.1038/s41586-018-0698-6 | https://www.cellph<br><br>onedb.org/ |
| msigdb 7.2.1 | Arthur Liberzon, Aravind Subramanian, Reid Pinchback, Helga Thorvaldsdóttir, Pablo Tamayo, Jill P. Mesirov, Molecular signatures database (MSigDB) 3.0, *Bioinformatics*, Volume 27, Issue 12, 15 June 2011, Pages 1739–1740, https://doi.org/10.1093/bioinformatics/btr260 | https://cran.r-<br><br>project.org/web/pa<br><br>ckages/msigdbr/in<br><br>dex.html |

| | | |
|---|---|---|
| DESeq2 1.32.0 | Love, M.I., Huber, W. & Anders, S. Moderated estimation of fold change and dispersion for RNA-seq data with DESeq2. Genome Biol 15, 550 (2014). https://doi.org/10.1186/s13059-014-0550-8 | https://bioconductor.org/packages/release/bioc/html/DESeq2.html |

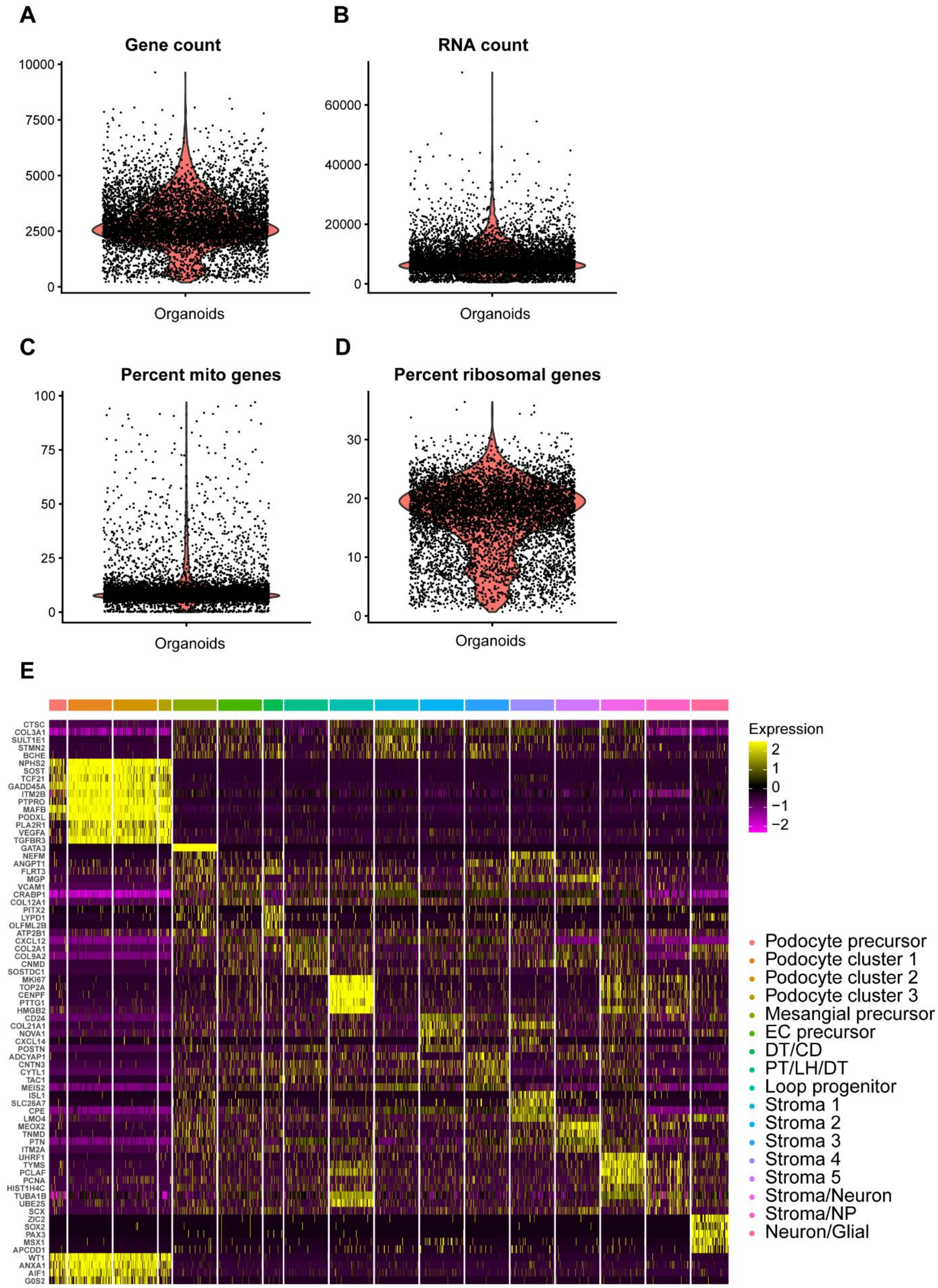


**Supplemental Figure S1, related to figure 1. Quality control data of scRNA seq data.** (A) Number of genes, (B) total number of RNA molecules, percentage of (C) mitochondrial and (D) ribosomal genes in the organoids sample. (E) Heatmap showing a selection of 5 out of the top 20 differentially expressed

(DE) genes per cluster. The full list of the top 20 DE genes per cluster is available in Supplemental Table S1. PT: proximal tubule, LH: loop of Henle, DT: distal tubule, CD: collecting duct, EC: endothelial cells.

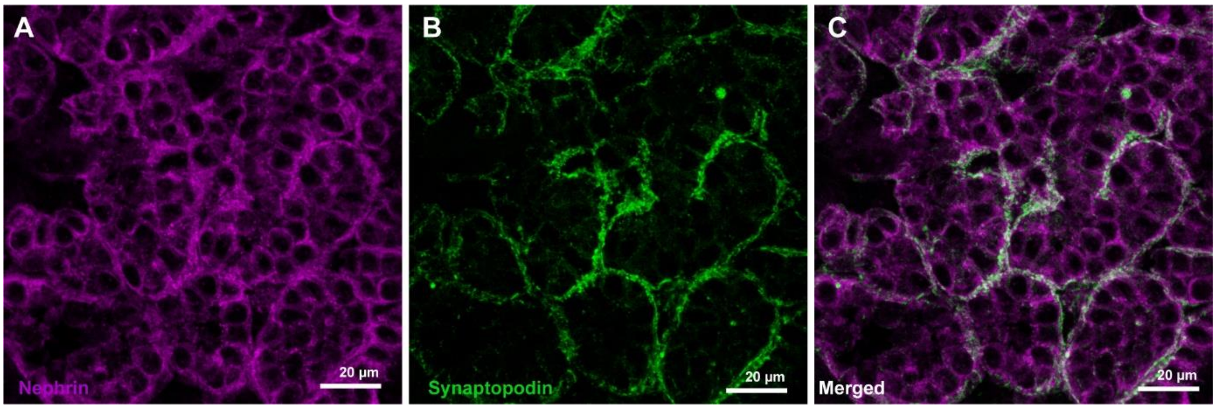


**Supplemental Figure S2, related to figure 1. Protein expression and localization of specific podocyte markers in kidney organoids.** (A) Nephrin (magenta) and (B) synaptopodin (green) expression in organoid podocytes (C, merged).

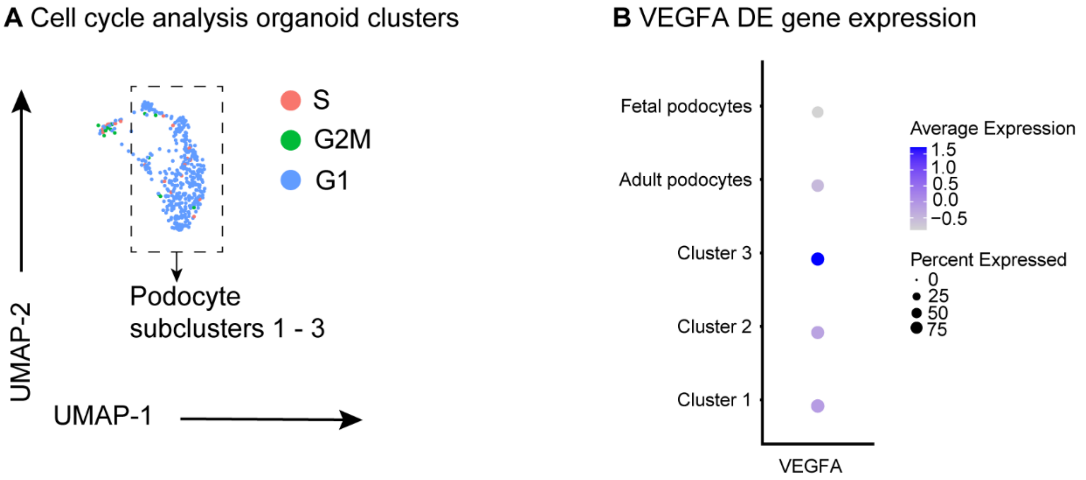


**Supplemental figure S3, related to figure 2. Cell cycle analysis of the podocyte sub-clusters and comparative VEGFA expression of organoid podocyte sub-clusters, human fetal and adult podocytes. DE: differentially expressed.**

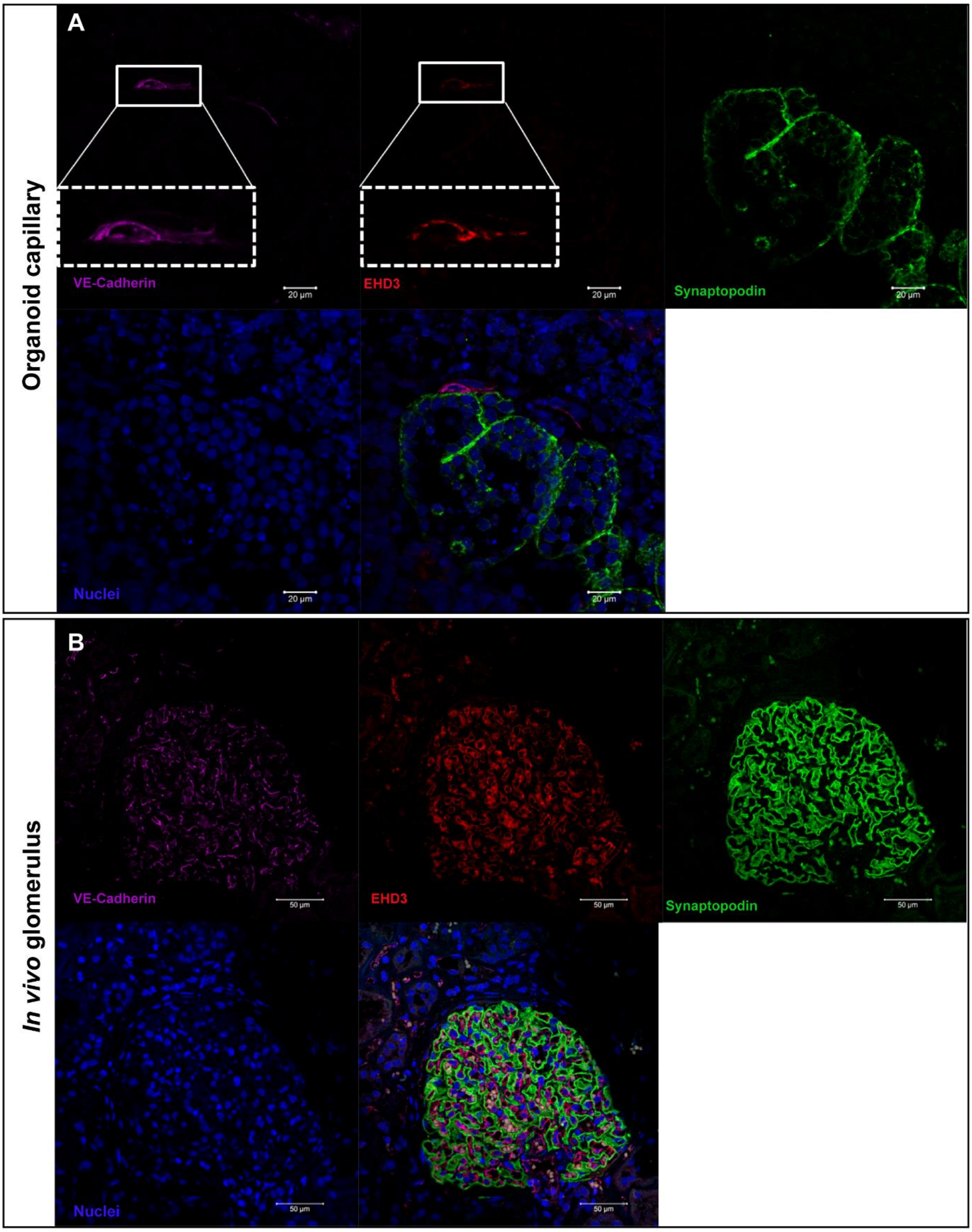


**Supplemental figure S4, related to figure 2. Glomerular endothelial EHD3 expression in organoids and in normal human glomeruli.** (A) VE-cadherin (magenta) and EHD3 (red) expression co-localized in an endothelial capillary in a kidney organoid, adjacent to synaptopodin expressing (green) podocytes. (B) VE-cadherin (magenta) and Eps15 Homology Domain-containing 3 (EHD3, red) expression co-localized in glomerular endothelium EHD3, in close proximity to podocytes as shown by synaptopodin expression.

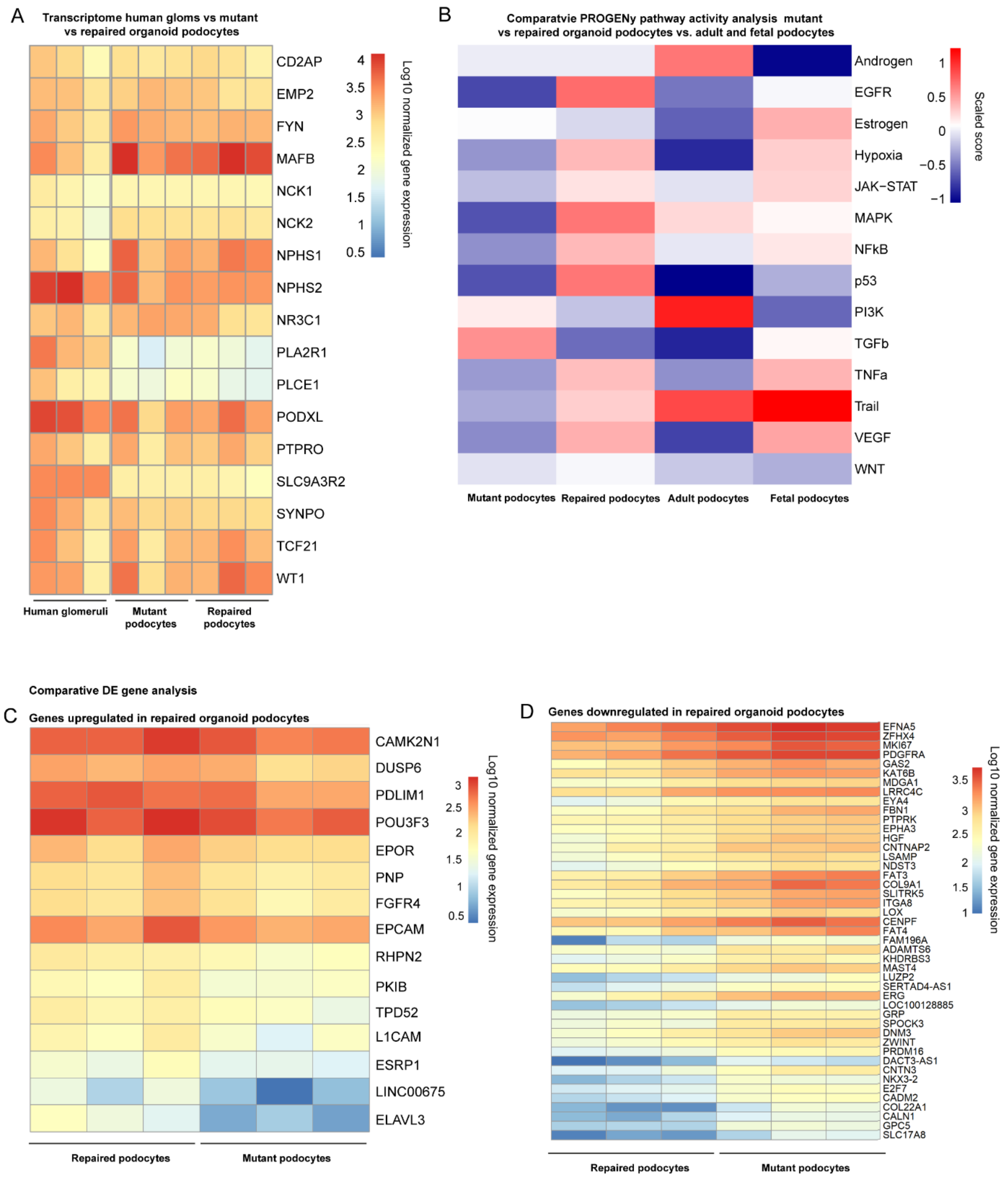


**Supplemental figure S5, related to figure 4. Comparative RNA sequencing analysis of kidney organoid *NPHS2* mutant podocytes, repaired organoid podocytes and normal human glomeruli-derived podocytes.** (A) Comparative bulk transcriptomics of podocyte specific genes in mutant and repaired organoid podocytes versus isolated human glomeruli. (B) Comparative PROGENy pathway activity analysis mutant and repaired organoid podocytes versus adult and fetal podocytes. Heatmap showing the enriched up- (C) and downregulated (D) genes in *NPHS2* repaired organoid-derived podocytes compared to mutant organoid-derived podocytes.

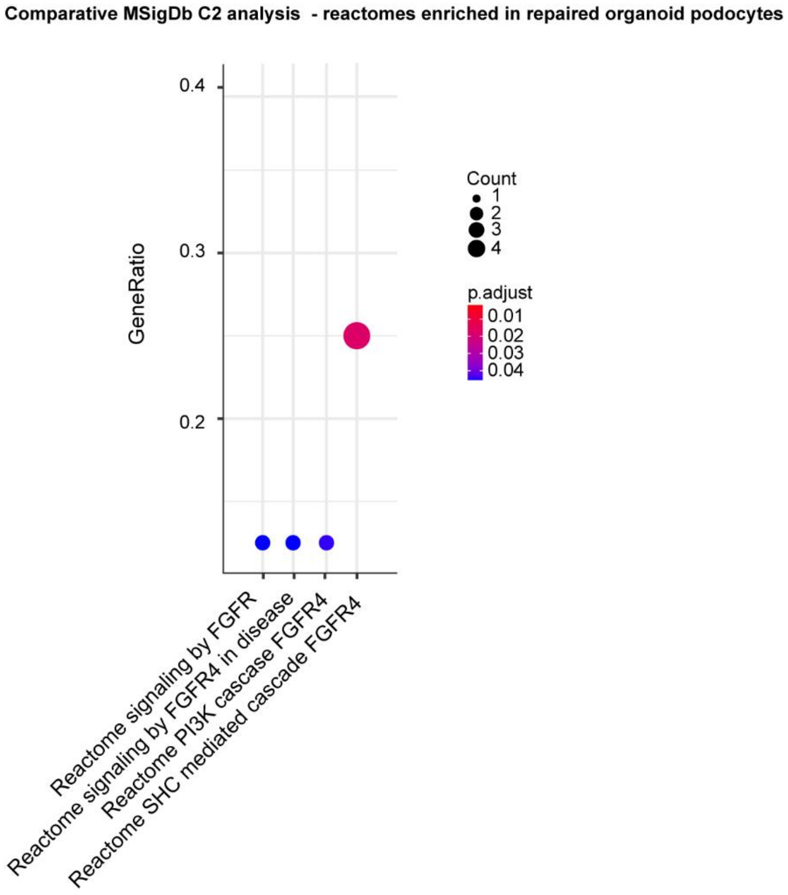


**Supplemental figure S6, related to figure 4. Comparative Molecular Signature database Analysis C2 showing enhanced FGFR reactomes in repaired organoid podocytes.**

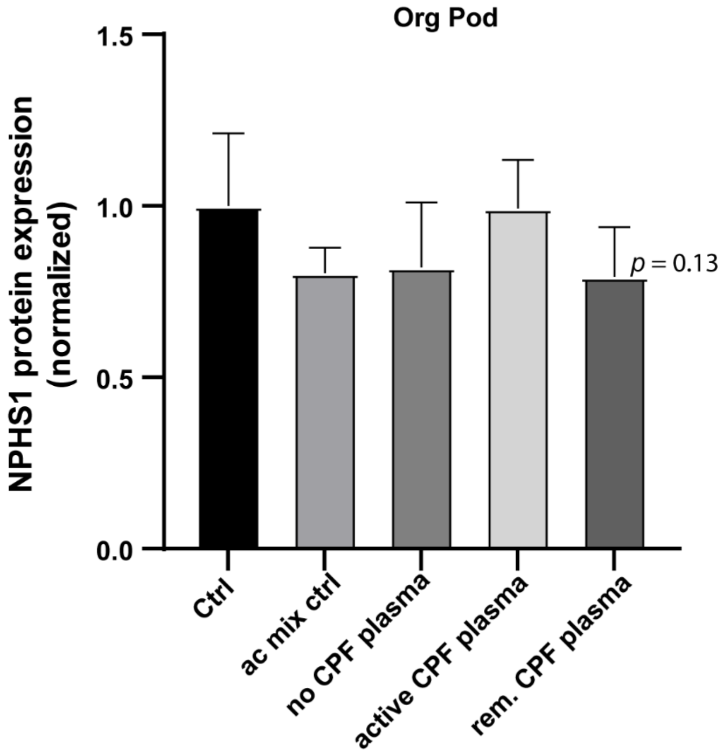


**Supplemental figure S7, related to figure 5. NPHS1 protein expression is not affected by FSGS plasma treatment.** Flow cytometry analysis of NPHS1 expression in organoid podocytes (Org Pod) treated with 10% (v/v) active and remission CPF plasma, CPF-free plasma, anticoagulation control (Ac mix ctrl) and E6 medium control (Control) for 4h.

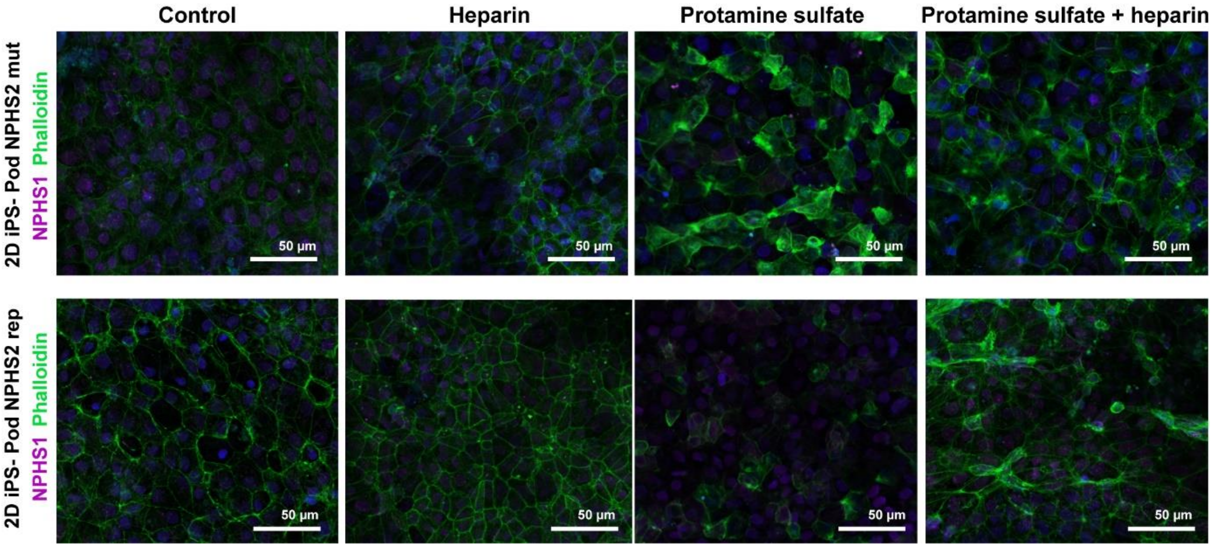


**Supplemental figure S8, related to figure 6. Cytoskeleton rearrangements in injured 2D iPSC-derived podocytes can be reversed upon heparin treatment.** Podocyte cytoskeleton analysis (phalloidin (green)) in mutant and repaired 2D iPSC-derived podocytes (NPHS1 (magenta)) following protamine sulfate treatment and heparin rescue.
